# On the Polymorph-Selection Determinants of α- Synuclein Amyloid Fibrils Studied at Atomic Resolution

**DOI:** 10.64898/2026.07.13.737774

**Authors:** Lukas Frey, David Rhyner, Witek Kwiatkowski, Dhiman Ghosh, Kristina Biedermann, Roland Riek, Jason Greenwald

**Author notes:** contributed equally to the work (Lukas Frey, David Rhyner).

## Abstract

Alpha-synuclein is an intensely studied intrinsically disordered protein whose aggregation into amyloid fibrils is connected to the progression of several neurodegenerative diseases, most commonly Parkinson’s Disease. A remarkable feature that has emerged from this research is how easy it is to induce the protein to aggregate *in vitro* into a wide range of amyloid fibrils that appear to resemble the aggregates found in Lewy bodies in diseases like Parkinson’s while at the same time how difficult it is to produce aggregates whose fold truly represents the disease-associated amyloids at the atomic level. In an effort to produce the disease-relevant fibrils *in vitro* we have analyzed over 60 independent samples by cryo-electron microscopy using helical reconstruction to obtain atomic resolution models for most of the samples. While not yet achieving our original goal, we have found that several overlooked parameters influence the structural outcomes of alpha-synuclein aggregation, in particular protein purity, preparation of the monomeric starting material and agitation method.

## INTRODUCTION

The capacity for structural polymorphism in amyloid fibrils has been recognized for decades but only recently has a large data set of atomic resolution amyloid structures become available, giving new insights into the nature of their multifaceted folds. This latter development is thanks largely to the power of cryo-electron microscopy-based helical reconstruction which has reduced the time and material needed to obtain a structure down to hours and micrograms. We have taken advantage of this technique to explore the determinants of polymorph selection in alpha-synuclein (α-Syn), a principle component of the Lewy body deposits associated with a number of neurodegenerative diseases that are collectively known as synucleinopathies (Goedert et al. 2017, Calabresi et al. 2023). While the degree of polymorphism is typically high in non-functional “disease-associated” amyloids α-Syn is perhaps the most polymorphic of all known amyloid-forming proteins: to date there are over 200 α-Syn amyloid structures in the PDB representing at least 15 different folds (depending on the fold classification scheme used). Like all non-functional amyloids, its protein sequence has not been optimized for any particular amyloid fold, giving rise to various mis-folding pathways that lie on a rough energy landscape with many minima in which a fold can be kinetically trapped. An extremely intriguing aspect of this pathway-dependent mis-folding lies in the correlation between the polymorph of the amyloids extracted from patients and the distinct pathological signatures of their disease (Schweighauser et al. 2020, Shi et al. 2021, Yang et al. 2022, Yang et al. 2023). That is, for the diseases belong within a single proteinopathy (i.e. synucleinopathies or tauopathies) the observed polymorph is both specific to that disease and challenging to reproduce *in vitro* (Lovestam et al. 2021). Because the in vitro polymorphs have been extensively explored at physiological conditions, there appears to be a selection pressure - a pathway and/or environmental influence on the polymorph outcome that occurs during the disease progression. Our previous results showed that there is a strong dependence on pH in the *in vitro* selection of α polymorphs with Type 1 being favored above pH 7 and Type 3 being favored below pH 6.8 while Type 2 appeared between pH 6.5-7.5 (Frey et al. 2024). We and others have reported on the secondary nucleation-dominated nature of α-Syn aggregation and how, unlike in elongation, the non-specific interactions involved do not preserve the seed polymorph (Peduzzo et al. 2020, Kumari et al. 2021). In an effort to produce the disease-relevant polymorphs of α- Syn *in vitro*, in particular the Parkinson’s disease (PD) polymorph, we have amassed a large dataset of amyloid structures (Table 1, Figure S1). Our results indicate that seemingly minor differences in sample preparation, the agitation method, and even trace protein impurities can completely alter the structural outcomes of α-Syn amyloid formation. We note that helical reconstruction inherently excludes non-twisted filaments, low-abundance polymorphs, and structurally heterogeneous species that fail to yield interpretable 2D classes. The polymorph outcomes reported here therefore represent a lower bound on the true structural complexity present in each sample.

**Table 1:** Samples for which Cryo-EM data was analyzed in this study. The outlines around sets of samples indicate those that were prepared on the same day and aggregated on the same rotator or same 96-well plate. Abbreviations: AMP=adenosine monophosphate, ADP= adenosine diphosphate, NADP= Nicotinamide adenine dinucleotide phosphate, PIP2= Phosphatidylinositol 4,5-bisphosphate PGE= Prostaglandin E_1_, DP = Dipyridamole, PT = Pemetrexed, rot^Q^ = samples during which the rotator stopped, PR^LB^ = low-binding 96- well plate, PR_UV_ = UV transparent 96-well plate, O-450 = orbital shaking at 450 rpm, ND = resolution too low to determine symmetry, NT = non-twisted or too few twisted for 3D analysis, HBS = 25 mM HEPES with 100 mM NaCl, PBS = phosphate buffered saline _1_ Samples **2G**-**36** are 4 pairs of samples from days 4 and 6 of 4 independent aggregations. _2_ Samples **56**+**57** are from days 8 and 12 of the same aggregation _3_ Samples **51**-**53** are Era 2 with 300 kD filter _4_ Sample **5G** has two distinct 1A polymorphs _5_ (5B) in parentheses indicates that it is clearly identified in micrographs but too few segments or irregular twist hindered further analysis _6_ The 1m fold variant in sample **17** is does not have the N-terminal strand (residues 10-17) _7_ The numbers in parentheses are FSC values without postprocessing for those datasets whose resolution was stuck at > 4.7 Å. _8_ The filled circles indicate that coordinates and/or maps have been deposited. Entry codes for deposited data are in Table S1

| sample | conc.<br>[$\mu\text{M}$] | buffer | pH | ligand /<br>mixture | sample<br>prep. | batch | purity | agitation<br>method | polymorphs observed | helical symmetry | postprocessed FSC<br>resolution<sup>7</sup> [$\text{\AA}$] | coordinates<br>deposited | maps<br>deposited |
| --- | --- | --- | --- | --- | --- | --- | --- | --- | --- | --- | --- | --- | --- |
| 1 | 300 | PBS | 7.0 | - | Era 1 | 1 | SP | rot | 1m, 2A, 2B | C1, C2, pC21 | (4.8), 2.8, 3.1 | O,O,O<sup>8</sup> | O,O,O |
| 2 | 300 | PBS | 7.0 | 600 $\mu\text{M}$ AMP | Era 1 | 2 | HPLC | rot | 5A | C2 | 3.03 | O | O |
| 3 | 300 | PBS | 7.0 | 600 $\mu\text{M}$ PGE | Era 1 | 2 | HPLC | rot | 5A, 5B | C2, pC21 | 2.97, 3.62 | O,O | O,O |
| 4 | 300 | PBS | 7.0 | - | Era 1 | 3 | SP | rot | 1m, 2A, 2B | C1, C2, pC21 | (4.8),(4.9),(4.9) | O,O,O | O,O,O |
| 5 | 300 | PBS | 6.8 | - | Era 1 | 4 | SP | rot<sup>Q</sup> | 3B, 3C | pC21, C2 | 2.6, 3.02 | O,O | O,O |
| 6 | 300 | PBS | 7.0 | - | Era 1 | 4 | SP | rot<sup>Q</sup> | 1A, 3B | pC21, pC21 | 2.8, 3.03 | O,O | O,O |
| 7 | 300 | PBS | 7.0 | - | Era 1 | 4 | SP | rot<sup>Q</sup> | 1A, 3B | pC21, pC21 | (6.5), (8.8) | O,O | O,O |
| 8 | 300 | PBS | 7.0 | - | Era 1 | 5 | SP | rot | 2A, 2B | C2, pC21 | 2.86, 2.95 | O,O | O,O |
| 9 | 300 | PBS | 7.0 | - | Era 1 | 6 | SP | rot | 1A | pC21 | 2.95 | O | O |
| 10 | 300 | PBS | 7.0 | - | Era 1 | 6 | SP | rot | NT |  |  |  |  |
| 11 | 300 | PBS | 7.0 | - | Era 1 | 6 | SP | rot | 1A | pC21 | 2.8 | O | O |
| 12 | 300 | PBS | 5.8 | - | Era 1 | 7 | SP | rot | 3B, 3C | pC21, pC21 | 3.94, 2.92 | O,● | O,● |
| 13 | 300 | PBS | 7.4 | - | Era 1 | 7 | SP | rot | -NT- |  |  |  |  |
| 14 | 300 | PBS | 7.0 | 2 mM AMP | Era 1 | 8 | SP | rot | 1m | C1 | 3.13 | O | O |
| 15 | 300 | PBS | 7.0 | 2 mM DP | Era 1 | 8 | SP | rot | 1m | C1 | 2.68 | ● | ● |
| 16 | 300 | PBS | 7.0 | 2 mM ADP | Era 1 | 8 | SP | rot | 1m | C1 | 2.83 | O | O |
| 17 | 300 | PBS | 7.0 | 1 mM PIP2 | Era 1 | 8 | SP | rot | 1m<sup>6</sup> | C1 | 3.25 | O | O |
| 18 | 300 | PBS | 7.0 | 2 mM NADP | Era 1 | 8 | SP | rot | 1m | C1 | 2.83 | O | O |
| 19 | 300 | PBS | 6.3 | - | Era 1 | 9 | SP | PR<sup>LB</sup> | 2E<sup>NS</sup> | pC21 | 2.63 | ● | ● |
| 20 | 300 | PBS | 6.3 | - | Era 1 | 9 | SP | PR<sup>LB</sup> | 2E<sup>NS</sup>, 3B, 3C | pC21, ND, pC21 | 3.21, (5.2), 3.13 | O,O,O | O,O,O |
| 21 | 300 | PBS | 6.3 | - | Era 1 | 9 | HPLC | PR<sup>LB</sup> | 2A, 3C | C2, pC21 | 2.89, 3.08 | O,O | O,O |
| 22 | 300 | PBS | 6.3 | - | Era 1 | 9 | HPLC | PR<sup>LB</sup> | 2A<sup>NS</sup>, 2B<sup>NS</sup> | C2, pC21 | 3.08, 3.05 | O,O | O,O |
| 23 | 300 | PBS | 7.0 | - | Era 1 | 9 | HPLC | PR<sup>LB</sup> | 2A, 2B, 5A, 5B | C2, pC21, C2, pC21 | 2.6, 2.86, 2.8, 2.66 | ●,●,●,● | ●,●,●,● |
| 24 | 300 | PBS | 7.0 | - | Era 1 | 9 | HPLC | PR<sup>LB</sup> | 2A, 5A, 5B | C2, C2, pC21 | 2.99, 2.83, 3.56 | O,O,O | O,●,O |
| 25 | 300 | PBS | 7.4 | - | Era 1 | 9 | SP | PR<sup>LB</sup> | 1m | C1 | 3.13 | O | O |
| 26 | 300 | PBS | 7.4 | - | Era 1 | 9 | SP | PR<sup>LB</sup> | 1m | C1 | 2.8 | O | O |
| 27 | 300 | PBS | 7.4 | - | Era 1 | 9 | HPLC | PR<sup>LB</sup> | 5m, 5B | C1, pC21 | 2.95, 3.2 | O,O | O,O |
| 28 | 300 | PBS | 7.4 | - | Era 1 | 9 | HPLC | PR<sup>LB</sup> | 5m, 5A, 5B | ND, ND, ND | (5.2), (4.7), (5.1) | O,O,O | O,O,O |
| 29<sup>1</sup> | 300 | PBS | 7.4 | - | Era 1 | 10 | SP | PR<sup>LB</sup> | 1m, 2A, 2B | C1, C2, ND | 3.06, (4.9), (5.4) | O,O,O | O,O,O |
| 30<sup>1</sup> | 300 | PBS | 7.4 | - | Era 1 | 10 | SP | PR<sup>LB</sup> | 1m, 2A, 2B | C1, C2, pC21 | 3.33, 3, 3.17 | O,O | O,O |
| 31<sup>1</sup> | 300 | PBS | 7.4 | - | Era 1 | 10 | SP | PR<sup>LB</sup> | 1m | C1 | 3.33 | O | O |
| 32<sup>1</sup> | 300 | PBS | 7.4 | - | Era 1 | 10 | SP | PR<sup>LB</sup> | 1m | C1 | 3.02 | O | O |
| 33<sup>1</sup> | 300 | PBS | 7.4 | - | Era 1 | 10 | SP | PR<sup>LB</sup> | 1m | C1 | 3.06 | O | O |
| 34<sup>1</sup> | 300 | PBS | 7.4 | - | Era 1 | 10 | SP | PR<sup>LB</sup> | 1m | C1 | 2.95 | O | O |
| 35<sup>1</sup> | 300 | PBS | 7.4 | - | Era 1 | 10 | SP | PR<sup>LB</sup> | 1m | C1 | 3.38 | O | O |
| 36<sup>1</sup> | 300 | PBS | 7.4 | - | Era 1 | 10 | SP | PR<sup>LB</sup> | 1m | C1 | 3.06 | O | O |
| 37 | 300 | PBS | 7.0 | 2 mM ADP | Era 1 | 10 | SP | PR<sup>LB</sup> | 1m | C1 | (4.9) | O | O |
| 38 | 300 | PBS | 7.0 | 2 mM AMP | Era 1 | 10 | SP | PR<sup>LB</sup> | 1m | C1 | 3.39 | O | O |
| 39 | 300 | PBS | 7.0 | 2 mM NADP | Era 1 | 10 | SP | PR<sup>LB</sup> | 1m | C1 | (4.8) | O | O |
| 40 | 300 | PBS | 7.0 | 2 mM PT | Era 1 | 10 | SP | PR<sup>LB</sup> | 1m | C1 | 3.29 | O | O |
| 41 | 100 | PBS | 7.0 | - | Era 1 | 11 | SP | PR<sup>LB</sup> | 1m | C1 | (4.6) | ○ | ○ |
| 42 | 100 | PBS | 7.4 | - | Era 1 | 11 | SP | PR<sup>LB</sup> | 1m | C1 | 2.71 | ○ | ○ |
| 43 | 100 | PBS | 7.4 | - | Era 1 | 11 | SP | PR<sup>LB</sup> | 1m | C1 | (4.9) | ○ | ○ |
| 44 | 300 | PBS | 7.4 | - | Era 1 | 12 | HPLC | O-450 | 2A, 6m, 6A | C2, C1, pC21 | 3.33, 3.21, 3.17 | ○,●,● | ○,●,● |
| 45 | 200 | PBS | 7.0 | - | Era 1 | 13 | HPLC | PR<sup>UV</sup> | 1A, 1/7, 2A, 5A, (5B)<sup>5</sup> | pC21, C1, C2, pC21 | 2.89, 2.68, 2.95, 2.86 | ●,●,●,● | ●,●,●,● |
| 46 | 200 | PBS | 7.0 | 5% NΔ4 | Era 1 | 13 | HPLC | PR<sup>UV</sup> | 1A | pC21 | 2.52 | ● | ● |
| 47 | 200 | PBS | 7.0 | 10% NΔ4 | Era 1 | 13 | HPLC | PR<sup>UV</sup> | NT |  |  |  |  |
| 48 | 200 | PBS | 7.0 | 50% NΔ4 | Era 1 | 13 | HPLC | PR<sup>UV</sup> | 1/7 | C1 | 2.2 | ● | ● |
| 49 | 200 | PBS | 7.0 | 100% NΔ4 | Era 1 | 13 | HPLC | PR<sup>UV</sup> | 8/9 | C1 | 2.65 | ● | ● |
| 50 | 200 | PBS | 7.0 | 100% NΔ4 | Era 1 | 13 | HPLC | PR<sup>UV</sup> | 8/9 | C1 | (6.9) | ○ | ○ |
| 51 | 150 | HBS | 7.0 | - | Era 2<sup>3</sup> | 14 | HPLC | PR<sup>UV</sup> | 2A<sup>NS</sup> | ND | (5.1) | ○ | ○ |
| 52 | 300 | HBS | 7.0 | - | Era 2<sup>3</sup> | 14 | HPLC | PR<sup>UV</sup> | 1m, 5m, 5A, (5B)<sup>5</sup> | C1, C1, C2 | 2.74, 2.77, 2.92 | ○,○,○ | ○,○,○ |
| 53 | 280 | PBS | 7.0 | - | Era 2<sup>3</sup> | 15 | HPLC | PR<sup>UV</sup> | 5m | C1 | (5.1) | ○ | ○ |
| 54 | 280 | PBS | 7.0 | - | Era 2 | 15 | HPLC | PR<sup>UV</sup> | 2A<sup>NS</sup>, 2B<sup>NS</sup> | C2, pC21 | 2.92, 3.82 | ○,○ | ○,○ |
| 55 | 280 | PBS | 7.0 | 5% NΔ4 | Era 2 | 15 | HPLC | PR<sup>UV</sup> | 2A, 2B | C2, pC21 | 2.92, (5.2) | ○,○ | ○,○ |
| 56<sup>2</sup> | 280 | PBS | 7.0 | 50% NΔ4 | Era 2 | 15 | HPLC | PR<sup>UV</sup> | 2A<sup>NS</sup>, 2B<sup>NS</sup> | C2, pC21 | 2.86, 2.92 | ●,○ | ●,○ |
| 57<sup>2</sup> | 280 | PBS | 7.0 | 50% NΔ4 | Era 2 | 15 | HPLC | PR<sup>UV</sup> | 2A<sup>NS</sup>, 2B<sup>NS</sup> | C2, pC21 | 3.17, 2.89 | ○,○ | ○,○ |
| 58 | 280 | PBS | 7.0 | 100% NΔ4 | Era 2 | 15 | HPLC | PR<sup>UV</sup> | 2B<sup>NS</sup> | pC21 | 2.32 | ● | ● |
| 59 | 200 | PBS | 7.4 | - | Era 2 | 16 | HPLC | PR<sup>UV</sup> | 1A, 1A<sup>4</sup>, (5B)<sup>5</sup> | pC21, pC21 | 2.83, 2.71 | ●,● | ●,● |
| 60 | 200 | PBS | 7.4 | 5% NΔ4 | Era 2 | 16 | HPLC | PR<sup>UV</sup> | 1m, 1D | C1, C1 | 2.89, 2.89 | ●,● | ●,● |
| 61 | 200 | PBS | 7.4 | 50% NΔ4 | Era 2 | 16 | HPLC | PR<sup>UV</sup> | 1A | pC21 | 2.11 | ○ | ● |
| 62 | 200 | PBS | 7.4 | 100% NΔ4 | Era 2 | 16 | HPLC | PR<sup>UV</sup> | 1A | pC21 | 2.26 | ● | ● |

## RESULTS and DISCUSSION

### Type 5 polymorph is inhibited by trace truncation products

While trying to understand the polymorph variability that we have observed between different α-Syn preparations, we turned to reverse-phase HPLC as an analytical tool to control for the effects of sample purity. The standard protocol (SP) that we use for α-Syn expression and purification which produces an N-terminally acetylated full-length protein is detailed in the methods section and is one of several published protocols that forgo affinity tags, relying instead on the unique properties of α-Syn and its periplasmic localization (Huang et al. 2005). Nevertheless, the SP routinely produces a “single band” purity as monitored by SDS-PAGE and so we were moderately surprised to detect significant levels of impurities in the HPLC chromatogram of this sample (red trace Figure 1A). We found that the purity between samples varied but for some like that depicted in Figure 1A it was as low as 75% based on the 220 nm absorbance. In order to minimize the effects of purity from one preparation to another, we added a reverse-phase purification step to the SP, yielding a protein that was >98% pure as assayed by analytical reverse phase chromatography (dashed line Figure 1A). As we began to work with the HPLC-pure α-Syn one significant difference quickly became apparent: the type 5 polymorph which we had only observed once among dozens of different aggregations in the pH 7.0-7.4 range was now appearing in every aggregation experiment in this pH range (Figure 1B). The Type 5 polymorph of α-Syn is unique in that it is both the only fold that encompasses the N-terminal methionine and is also the largest fold, comprising up to 97 of the 140 residues in addition to around 14 residues whose identities remain unclear but are likely derived from the last 30 acidic C-terminal residues. The unique presence of the N-terminus in the fold as well as the possible involvement of residues at or near the C-terminus suggested that only a full-length α-Syn would be competent to form the type 5 polymorph. In particular, the interface of the 5A fold appears to require the acetylation of the N-terminal Met, as its methyl forms part of the interface with Val40 on the other chain (Figure 2B). To date, 5A has only been observed with acetylated α-Syn while other interface variants of type 5 have been published with α-Syn bearing a free amine at the N-terminus. This led us to look more closely at the potential contaminants in the SP protein and one preparation in particular that was used in aggregation batches 8-11 stood out because it consistently produced pure monofilament Type 1m polymorphs. This polymorph consistency spanned several different aggregation experiments at pH 7 and 7.4, many in the presence of a variety of small molecules that had been included in the hopes that they would encourage the formation of the PD-polymorph (Table1, samples **14**-**18**, **25**-**26**, **29**-**43**). We therefore performed high resolution liquid chromatography-mass spectrometry (LC-MS) analysis on the 25k g pellet and supernatant of an aggregation that had produced only type 1m. The pellet contained dozens of N- and C-terminally truncated species that eluted before the full-length α-Syn, the most abundant of which had signal intensities around 10% of the intact full-length protein (Figure S2). In contrast, only a single degradation product, Ac-(1-72), was observed in the supernatant but it had a relatively higher MS signal than in the pellet (20% versus 10% relative to the full-length) and the degree of oxidation of the full-length protein was also higher based on the relative signal intensity (28% versus 3%). A simple interpretation of these observations is that many of the degradation products are more aggregation prone than the full-length α-Syn and that the oxidation of a single methionine is sufficient to reduce the aggregation. However, it is also very likely that many of the degradation products resulted from the conditions under which the α-Syn was aggregated into the Type 1m amyloid: 5 days of agitation at 37° C. With this in mind, we selected the N-terminal truncation of the first four residues (NΔ4) out of the major degradation products in the pellet fraction for further study. Not only is NΔ4 interesting due to its reported presence in the SDS-insoluble fraction of human brain tissues (Kellie et al. 2014), it is also the closest in mass to the full-length protein and therefore could easily have escaped notice by SDS-PAGE even at non-trace levels. Since the fifth residue of α-Syn happens to be a methionine, we could also express the exact truncation product without any additional tags and purify it using the same SP method. A final HPLC purification step was included for the NΔ4 production in order to be certain that no other truncation products were present. The elution time of the HPLC-pure NΔ4 on an analytical HPLC column was similar to one of the impurities in the SP-α-Syn (Figure 1A) however, the LC-MS analysis of the purified NΔ4 sample revealed an extra methyl mass of 14.05 Da present in 40%- 70% of the sample (the amount varied from one expression to the next). At first thinking that there was a Val to Leu/Ile mutation, we sequenced the expression vector which revealed no errors and also ordered a new codon-optimized gene for *E. coli* expression in a different vector. Upon observing the same extra mass in subsequent expressions using the re-synthesized gene, we then had the sample analyzed by Endo Glu-C digestion followed by MS-MS analysis to determine that the extra mass was due to a methylation on the N-terminal Met residue. Also unexpected was the presence of low levels of methylation on a few Glu and Lys residues throughout the sequence (Figure S3). Since two independent expressions of the NΔ4 contained the methylation we decided to carry on with the analysis of its aggregation with the caveat that all NΔ4 samples contain a mixture of un-modified and N-terminally methylated methionine and that this modification had an unknown effect on the protein aggregation behavior. The HPLC-pure NΔ4 as well as its 5% and 50% mixtures with HPLC-pure α-Syn were then prepared to test the effect of this truncation on the aggregation outcomes (Table 1, samples **45**-**50**). As expected the pure NΔ4 does not form the type 5 polymorph, but more surprising is that at as low as 5% it seems to completely inhibit type 5 polymorph formation (Figure 2C). The 100% HPLC-pure α-Syn (the same stock used to make the mixtures with NΔ4) recapitulated the type 5A as well as the 1A, 2A and a novel asymmetric 1/7 two-filament fold (Figure 2D). The 100% NΔ4 produced another novel asymmetric two-filament fold which we have named type 8/9 (Figure 2C and 2D). In summary, the type 5 polymorph requires an intact N-terminus and its formation is inhibited by trace truncations such as the NΔ4 variant typically present in non-HPLC purified material.

**Figure 1:**
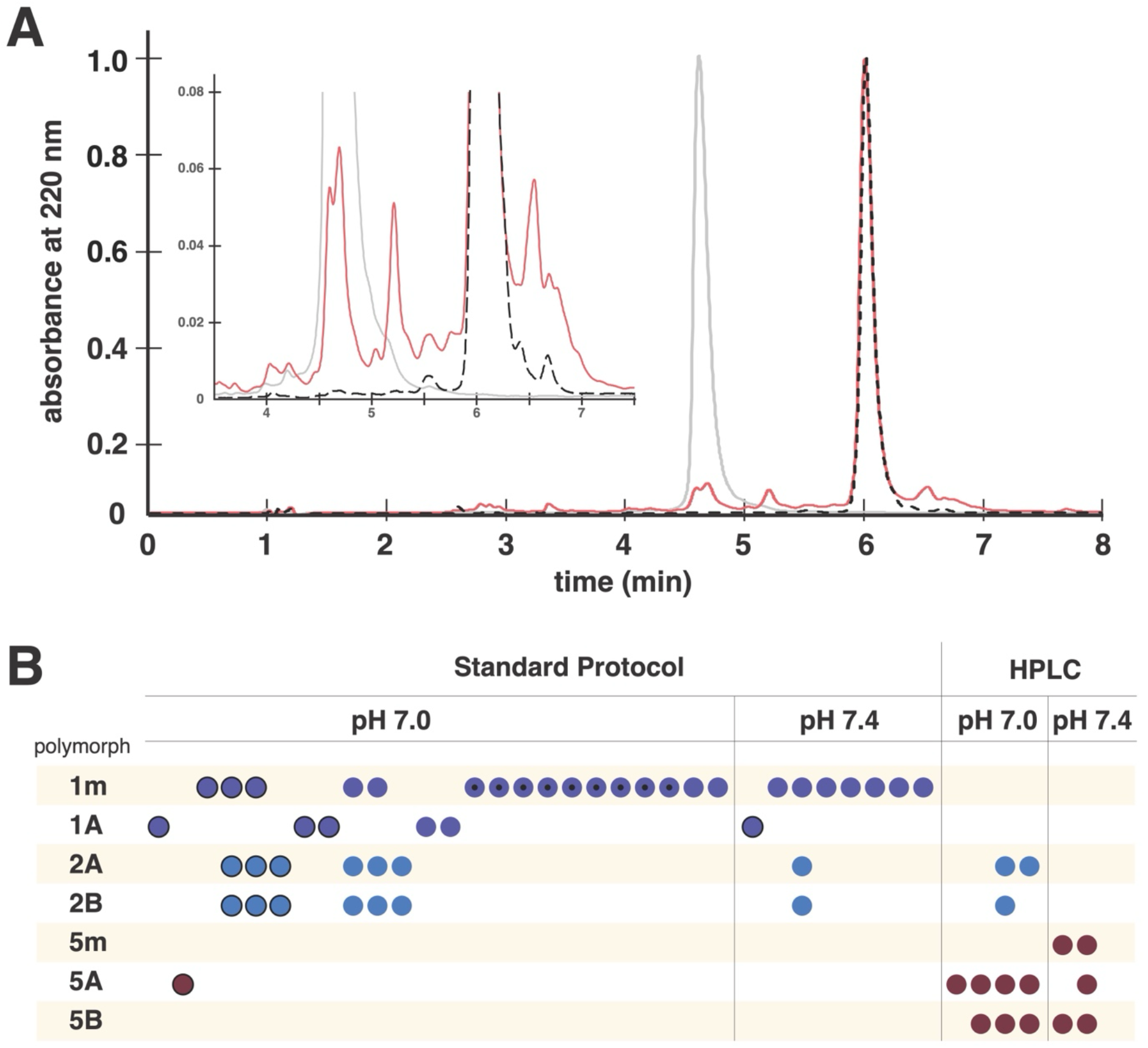
Purity of α-Syn is correlated with amyloid polymorph formation. (**A**) Analytical HPLC traces of SP-pure (red) HPLC-pure (black dashed) and NΔ4 α-Syn (grey) with inlay zoomed in on region around the main peak. (**B**) Dot plot of the polymorph outcomes for 38 independent aggregation experiments at pH 7.0 or 7.4 with α-Syn prepared with either the standard protocol or with additional HPLC purification. Each column represents one experiment with its Cryo-EM-verified polymorph outcome represented by a colored dot: purple = type 1, blue = type 2 and maroon = type 5. A black circle around the dot refers to samples that were previously reported in (REF) and those with a black point in the middle are samples for which a small molecule additive was included as part of a binding screen.

**Figure 2:**
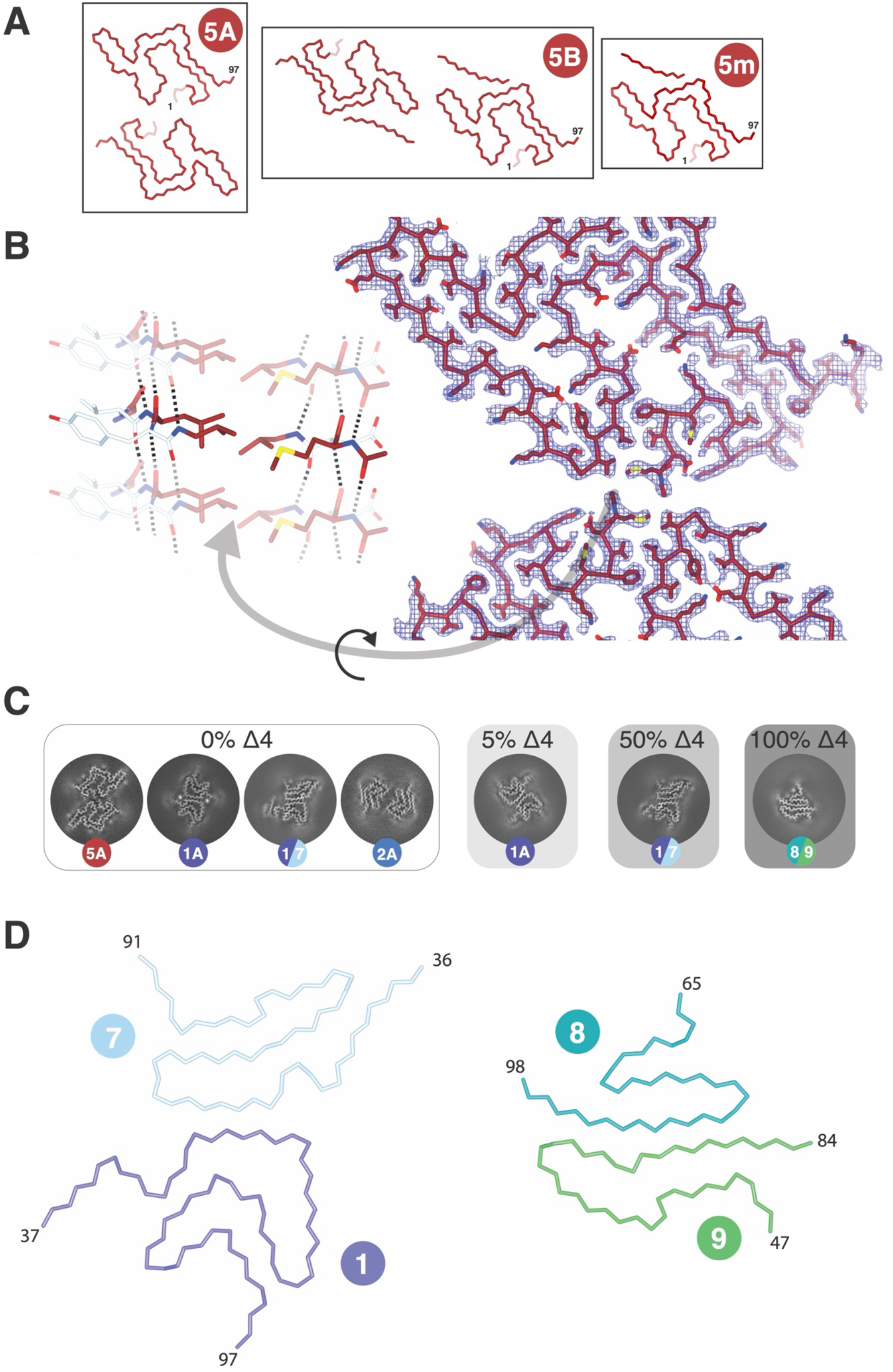
The type 5 polymorph requires an intact N-terminus and is inhibited by trace truncations. (**A**) The Cα traces of the type 5 assembly variants 5A, 5B and 5m showing the extent of the sequence coverage from residues 1-97 and, in the case of some 5A (not shown here) and for all 5B and 5m, an additional detached β-strand whose identity could not be determined. The first 4 residues which are truncated in the NΔ4 variant are highlighted to illustrate their location in the fold. (**B**) On the right, a view down the fibril axis of the Type 5A interface showing Cα trace with sidechains along with its EM density illustrates the extent of the hydrophobic interactions between Met1, Val3, Leu38, Val40 within the fold and for the N-acetylation at the 5A interface. On the left, a view rotated by 90° depicts three layers on one half of the 5A interface with the middle layer highlighted. For clarity, only the four residues mentioned above are colored while others are white. Dotted lines indicate the H-bonds between backbone atoms as well as those for the amide of the acetylated N-terminal methionine. (**C**) Z-slices (∼5 Å thick and 150 Å in diameter) for each Cryo-EM reconstruction found in the aggregated samples composed of 0%, 5%, 50% and 100% NΔ4. The colored dots are labeled with the polymorph types. (**D**) The two novel asymmetric polymorphs 1/7 and 8/9 depicted as Cα traces color coded by polymorph class and with the terminal residues of each chain indicated.

### Polymorph co-existence

It is important to note that the above mentioned study on the mixtures of full-length and NΔ4 protein, only the 100% HPLC-pure full-length α-Syn (sample **45**) produced a type 5 polymorph, but it is equally interesting to note that it was produced along with types 1A, 2A and a novel type 1/7 two-filament fold (Figure 2D). We could even identify single micrographs in sample **45** in which all four fibril types were present within an area of 0.1 μm_2_ (Figure 3A). In fact, there were at least three additional polymorphs present which could not be analyzed at high resolution due to their low population or low data quality, although one of them appeared to be 5B based on its unique 2D profile. Actually, the presence of multiple distinct α-Syn polymorphs within a sample is not so uncommon (Table 1) and considering that the nucleated growth and the likely difference in kinetics by which the various folds increase in number would be expected to bias the aggregation outcomes towards a single winner this polymorph co-existence is an interesting feature of the aggregation pathway. However, as demonstrated for Tau, aggregation pathways can be much more complicated than can be ascertained from ThT kinetics alone (Lovestam et al. 2024). Sample **23**, another HPLC pure sample, also displayed four co- existing polymorphs, this time with a different constellation of polymorphs, and they too were sometimes localized on a single micrograph (Figure 3B). In total, we observed 12 instances of polymorph class coexistence (not counting coexisting interface variants). Our feeling is that the occurrence of co-existing polymorphs is likely underreported in the literature due to selection biases in Cryo-EM which tend to exclude low population or low quality data in general. Also, the inherent challenges associated with some polymorphs, like low or irregular twist, often stalls their analysis at the 2D classification stage. In summary, polymorph co-existence for α-Syn amyloids is common even within femtoliter-scale volumes, indicating that identical local solution conditions can give rise to multiple fibril architectures rather than a single deterministic structural outcome.

**Figure 3.**
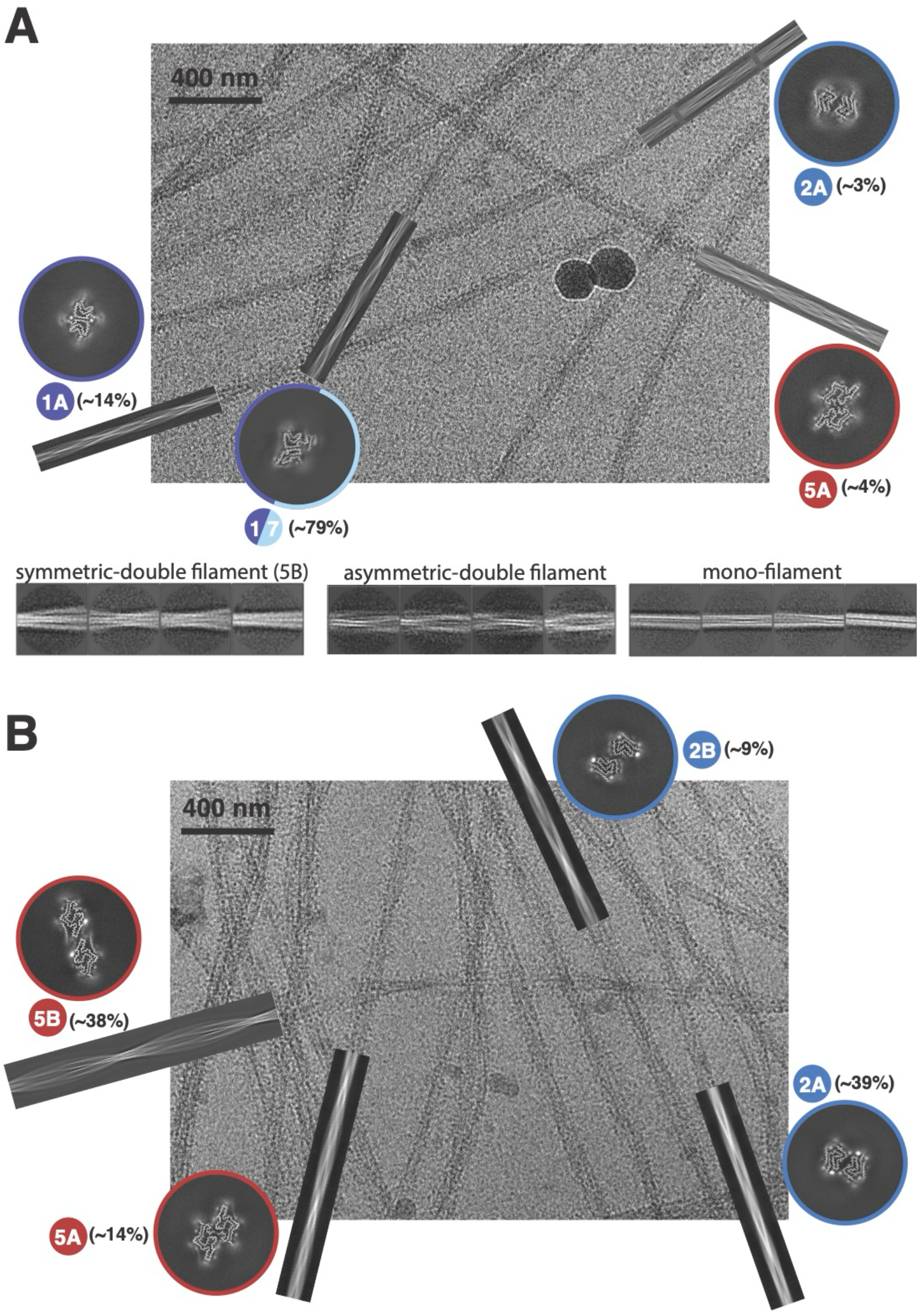
Multiple coexisting polymorphs appear on single micrographs. (**A**) A single micrograph from the HPLC-pure α-Syn sample from the Era 1 pH 7.0 (sample **45**) overlayed with projections of the initial 3D-models from relion_helix_inimodel2d aligned to fibrils of the same polymorph. The z-slices from the final refined 3D maps are ∼5 Å thick and labelled with the approximate relative amount of that polymorph in the dataset. Below the micrograph are three sets of 2D classes from the same dataset representing three additional fibril polymorphs that were not further analyzed due to their low abundance. (**B**) A single micrograph from an Era 1 pH 7.0 HPLC-pure α-Syn sample overlayed with the identified coexisting polymorphs, as in **A**.

### Agitation conditions can bias polymorph selection

In the course of screening new conditions for α-Syn aggregation we once tried a continuous vortex-like agitation at 450 rpm instead of our standard rotation or plate reader agitation protocols (see methods). An HPLC pure α-Syn sample at pH 7.4 under this strong agitation (sample **44**) did not lead to type 5 but rather to a mixture of the type 6m, 6B and 2A polymorphs (Figure 4A and 4B). This is surprising because the type 6 polymorph has only been observed once in the published literature (as 6m and 6A) and that was with an N-actetyl glucosamine (GlcNAc) addition on residue Ser87 (8JEX, EMD- 36202 (Hu et al. 2024)). In light of these published results, it is noteworthy that Ser87 is not visible in the 6m and 6B structures presented here. Nonetheless, their core type 6 folds are nearly identical and the published GlcNAc-modified 6A and the current 6B structures share some unusual characteristics. First, there is their distinct inter-filament interfaces which are both more than 20 Å wide and lack any visible contacts despite having some unidentified weak density between the filaments. Second, the 6A,6B as well as 6m fibrils all have a right-handed twist which is unusual for α-Syn amyloids. One notable exception is the elusive PD-polymorph which usually forms non-twisted fibrils but when it does form twisted fibrils they are also right-handed ((Yang et al. 2022)). The authors of the 6A structure concluded that the modified Ser87 was responsible for the novel polymorph but our result points to other explanations and highlights the difficulty in interpreting the structural outcomes of α-Syn aggregation. In another fortuitous discovery, we found that using less than our standard amount of agitation can also impact polymorph selection. For our previous publication on α-Syn (Frey et al. 2024) and for many of the samples in the current data, we used a lab rotator with a 3-4 second period to constantly but gently agitate 500 μl samples end over end in a microcentrifuge tube. During the aggregation of one set of 3 samples at pH 6.8 and 7.0 the rotator motor died sometime in the first 48 hours and the samples remained quiescent for the remainder of the aggregation (samples **5**-**7**). To our surprise, the pH 7.0 semi-quiescent aggregations both led to a mixture of type 1A and 3B (Figure 4A). Until now we had never observed these two polymorphs in the same sample because the pH ranges that favor their formation do not overlap. In the pH 6.8 semi-quiescent sample only 3B and 3C were observed which is also remarkable due to this pH usually favoring type 2 polymorphs. A new sample (**8**) was prepared from the same α-Syn prep and aggregated on the repaired rotator for the full 5-days of aggregation leading to the expected type 2A/2B polymorphs. These examples of variations in agitation leading to new outcomes in polymorph selection are in contrast to the lack of an observable effect between other agitation protocols we have used: the slow rotation of 500 μl samples and the intermittent orbital shaking of 200 μl samples in a 96-well plate (see Table 1 for all conditions and outcomes). The latter method was used for much of the data in this manuscript and allowed us to minimize on protein amount and also to monitor the kinetics of the aggregation (see methods). In summary, the agitation protocol should be added to the list of factors that can affect the polymorph outcome of α-Syn.

**Figure 4:**
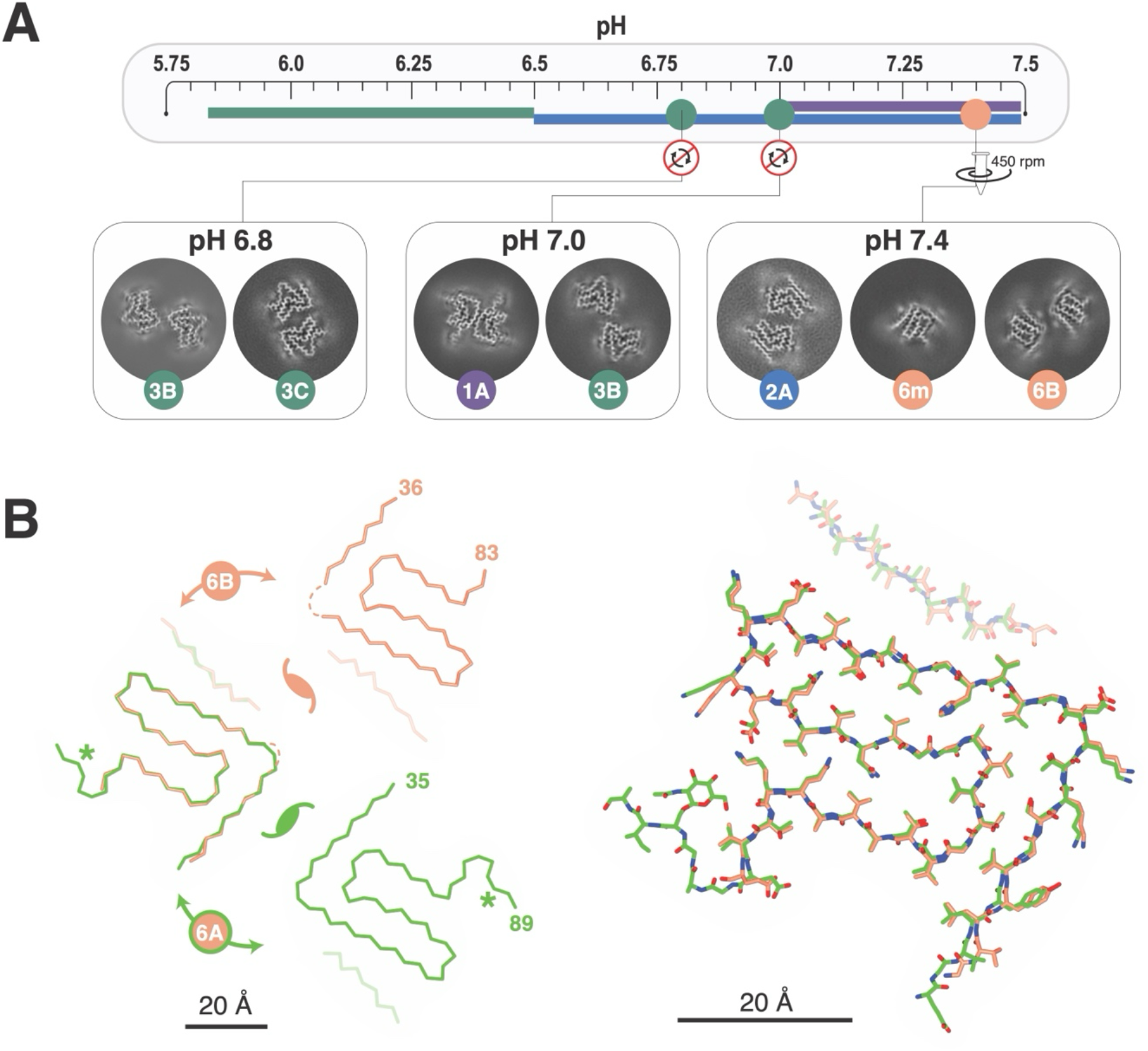
Agitation protocol influences polymorph outcomes. (**A**) The color-coded regions below the pH scale indicate the polymorph types that are preferred at each pH range (purple = 1, blue = 2, green = 3 ) as described in (Frey et al. 2024). The three colored circles on the pH scale indicate the occurrence of polymorphs under alternative agitation regimes and are connected by lines to their respective Cryo-EM map z-slices. The z-slices (∼5 Å thick and 150 Å in diameter) of the multiple coexisting polymorphs are grouped for each sample with the classes indicated in the color-coded circles. (**B**) Structural comparison of the type 6A (green carbon atoms) and 6B (orange carbon atoms) assembly variants. On the left, an overlay of a single subunit from each assembly variant highlights their distinct interfaces and unusually large inter-filament distances. The locations of their respective pseudo-C21 axes are indicated as are the positions of the N- and C-terminal residues. The position of the GlcNAc-modified Ser87 is indicated with the “*”. On the right is a superposition of a single unit of the 6A (pdb: 8JEX) and 6B showing the high degree of similarity, including the position of the additional isolated but unidentified strand (here shown in faded colors).

### Under reduced-oligomer sample preparation conditions, pH reasserts as the dominant polymorph selector

Our previous work outlined a set of pH rules for α-Syn polymorph selection while also pointing out the stochastic nature of the aggregation processes and the potential influence of unknown or uncontrollable parameters. In an effort to simplify sample preparation and provide starting material with the least influence from preformed seeds we modified our monomer sample preparation protocol for subsequent analyses of the HPLC-pure full-length and NΔ4 α-Syn aggregation. The ideal monomeric sample preparation is protein as it elutes from a size-exclusion column, however in practice this is not ideal because the concentration at which the protein elutes is difficult to control and typically too dilute for aggregation reactions. Our standard sample preparation (referred to as “Era 1”) involved the solubilization of α-Syn in the buffer of choice, adjusting the pH to 7-8 with NaOH, passing it over a PD-10 buffer exchange column and then concentrating the sample with a 10 kD MWCO ultrafiltration device before finally filtering the sample though a 100 kD MWCO ultrafiltration device. There were losses at several steps but particularly at the final filtration step which often removed 30-40% of the sample. The new “Era 2” sample preparation took advantage of the high solubility of lyophilized HPLC-purified α-Syn in pure water. This high solubility was probably a result of the pH in water falling well below the pI of α-Syn due to the trace TFA from the HPLC solvent that is not removed during lyophilization. However, when this sample was then dialyzed against the aggregation buffer to a pH above the pI, nearly 100% of the sample remained soluble. This way, the protein concentration could be maintained above the 200-300 μM that we typically used for aggregation reactions with minimal sample loss and avoiding the need for the concentration steps that are prone to aggregate formation. Then, we passed the protein over a 300 kD followed by a 100 kD MWCO filter (with the exception of samples **51**-**53** which were only passed over the 300 kD filter). We found that using two sequential filters prevented filter blockage and reduced the sample loss which was now only about 10%. The resulting “Era 2” HPLC-pure full-length and NΔ4 α-Syn samples and their mixtures yielded a very consistent pH dependence to the polymorph outcomes. At pH 7.0, the 0%, 5%, 50% and 100% NΔ4 mixtures (samples **54**-**58**) all yielded type 2 polymorphs and at pH 7.4 the same mixtures (samples **59**-**60**) all yielded type 1 polymorphs (Figure 5) including a new interface variant type 1D (Figure S4A and S4B). The consistency of the main class within samples at each pH is remarkable and aligns well with the original pH-polymorph trend that we found in the Era 1 samples in this and our previous work ((Frey et al. 2024)), further strengthening the pH-polymorph correlation. There was, however, still an assortment of variants within the type 1 and 2 classes from mixture to mixture at a given pH, discussed further below.

**Figure 5:**
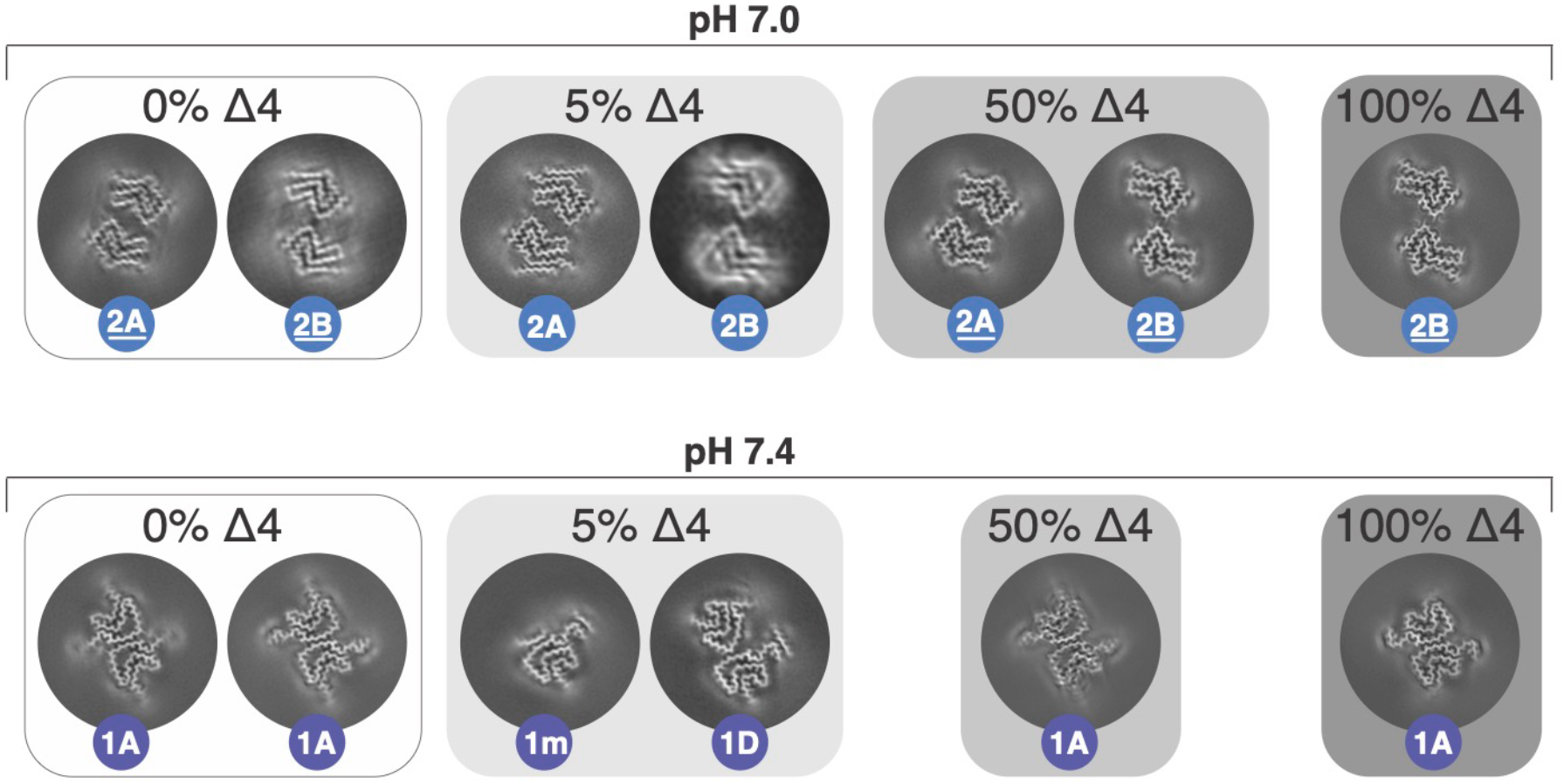
Under reduced oligomer conditions pH reasserts as dominant polymorph selector. The aggregation outcomes for HPLC-pure full-length and NΔ4 samples prepared under “Era 2” preparation conditions are shown as z-slices of the Cryo-EM maps (∼5 Å thick and 150 Å in diameter) with numbered color-coded balls to indicate the polymorph. The background shading of each aggregation experiment is indicative of the NΔ4%. The names of the type 2^-NS^ variants that lack the N-terminal strand are underlined.

Returning to the original motivation for the NΔ4 mixtures, the absence of type 5 from the HPLC-pure full-length sample in these Era 2 samples is noteworthy. However a visual inspection of all of the pH 7.4 micrographs from the NΔ4 series identified fibrils with the distinct Type 5B morphology only in sample **59**, the one without NΔ4 (Figure S5). The presence of a 5B polymorph would be consistent with the data presented so far, however the low frequency of these 5B-like fibrils precluded a 3D alignment of the segments. Nonetheless, polymorph 5 does appear in Era 2 preparations of pure full-length α-Syn for which the 100 kDa MWCO filter was omitted (see kinetics below).

In summary, an optimized monomer preparation protocol of HPLC purified α-Syn that aimed to avoid potential oligomer-forming steps produced polymorphs of single type per sample for which the pH appeared to be dominant polymorph selector.

### The primary nucleation-dependence of polymorph outcomes is indicated in aggregation kinetics

The idea that it is the absence or very low level of oligomer species (or perhaps amyloid seeds) that allows pH to become the dominant polymorph determinant is supported by two other Era 2-like preparations for which we only carried out the 300 kD filtering step, leaving out the 100 kD filter step. The first, sample **53** was from the same pH 7.0 preparation that led to the type 2 structures but was derived from a portion of the sample that only passed over the 300 kD filter and was also allowed to aggregate on the same 96- well plate. The second, sample **52**, was a part of another small molecule screen for which we needed to replace the phosphate with another buffer so we used HEPES buffered saline at pH 7.0, also only passed through the 300 kD filter. Both of these samples produced type 5 polymorphs and since the aggregation kinetics of all Era 2 samples was followed simultaneously by turbidity (A340 nm) and by tyrosine fluorescence polarization (FP) (Figure 6) we could gain some insight into the differences in kinetics for the 100 kD versus 300 kD filtered samples. These A340 and FP measurements offered complementary observations of the onset of aggregation, both having different sensitivities to the size and flocculence of the aggregate and possibly to the polymorph because the dynamic flexibility of the tyrosine residues should impact FP but not A340. An assessment of the aggregation kinetics compared to the polymorph outcomes suggests that faster aggregation onset (shorter lag time) in otherwise similar conditions are more likely to produce Type 5 and longer times are more likely to have pH-selected outcomes. The HEPES pH 7.0 sample **52** mentioned above (300 kD-filtered Era 2-like) was one of three samples in a concentration series (300 μM, 150 μM and 75 μM) for which the lower concentrations were made as 2x and 4x dilutions in filtered buffer. The aggregation kinetics of these three samples is shown in Figure 6A along with the observed polymorph outcome for two of the samples. Although the aggregation kinetics profiles were not fit to any amyloid assembly models, the curves are indicative of very different processes and different outcomes. Secondary nucleation seems more dominant in the 75 μM and 300 μM sample while the 150 μM sample appears to be a slower process whose rate may be limited by the addition to the ends of the growing filaments. The lower A340 signal for the 75 μM sample is expected due to its lower concentration but its higher FP signal suggests that it may have yielded a higher percent of aggregate or formed an amyloid in which its tyrosine residues are less flexible. The appearance of the types 5m and 5A in sample **52** with the shorter lag-time (300 μM) also suggests that that fold was derived from seeds or an aggregation competent form of α-Syn that was absent or below a critical concentration in the 150 μM sample **51** in which the type 2A fibrils were formed. Likewise, sample **53**, the other 300 kD-filtered sample of HPLC-pure full-length α-Syn from the Era 2 NΔ4 mixtures at pH 7.0 displayed completely different aggregation kinetics (yielding type 5m) and had a significantly shorter lag time compared to the same sample that had been passed over a 100 kD filter (yielding types 2A_-NS_ and 2B_-NS_) (Figure 6C). The molecular mass of monomeric α-Syn is near 14 kD meaning that even with the 100 kD filter some oligomeric forms could pass through into the filtrate, however, using a smaller filter is not possible because the unfolded nature of α-Syn makes its effective size in solution more like a protein of 3-4 times its mass which would lead to large losses in a filter with a smaller cutoff. The 300 kD filter on the other hand appears to be too large to prevent some seeding competent forms of the protein though, which may be in the range of a 10-20mer but still significantly smaller than the type of seeds produced from mature amyloids though sonication or other mechanical methods for seeding experiments.

**Figure 6:**
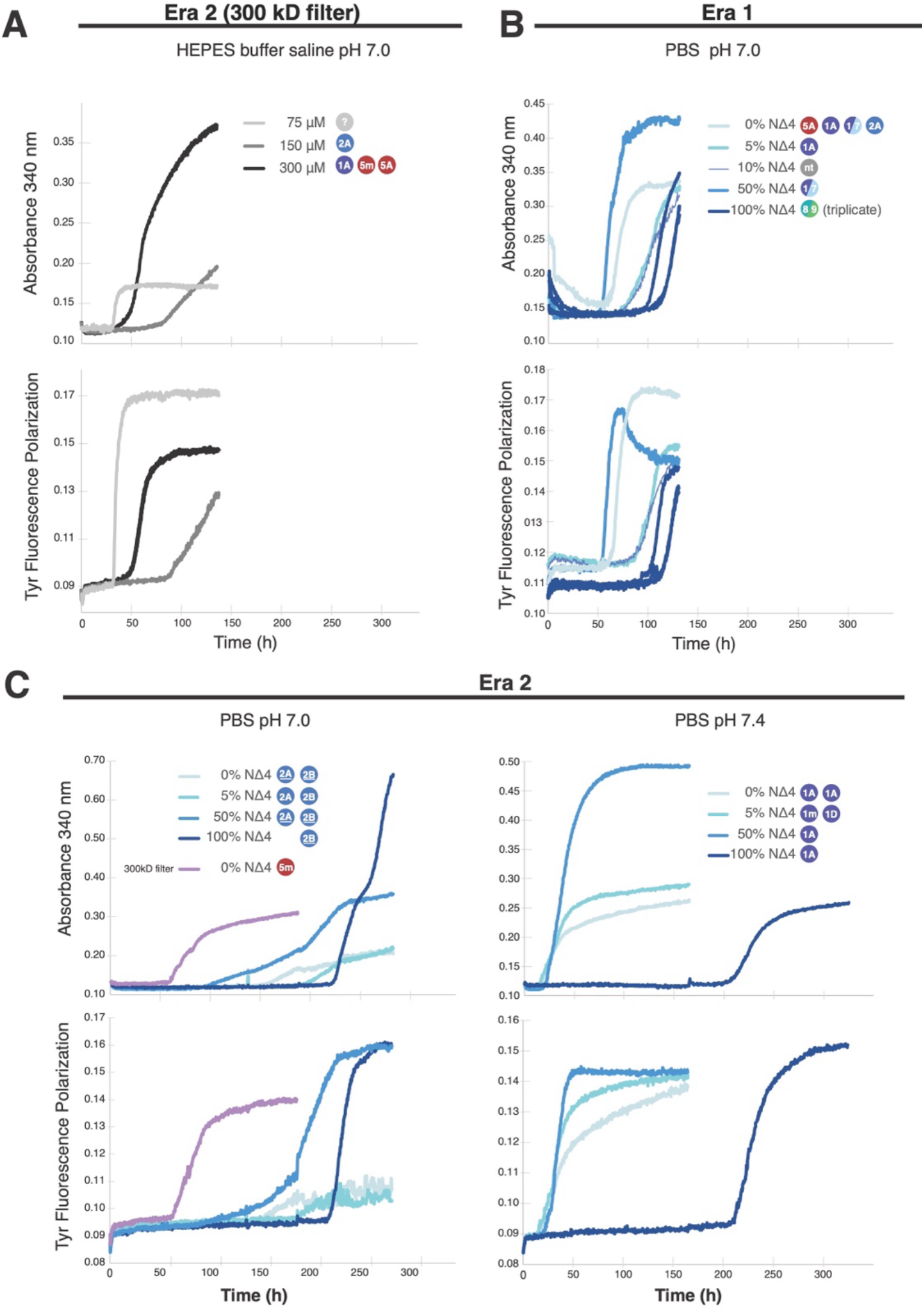
Correlating aggregation kinetics to polymorph outcomes hints at seed-dependence. (**A**) The aggregation kinetics of three independent HPLC-pure α-Syn samples followed by simultaneous measurement of absorbance at 340 nm (upper plot) and tyrosine fluorescence polarization (lower plot). The three samples were prepared by serial dilutions from a single 300 μM stock which had only been passed over a 300 kD filter (no 100 kD filter). The polymorph outcomes of the 300 μM and 150 μM aggregations (samples **51** and **52**) were analyzed by Cryo-EM and noted as colored balls. (**B**) and (**C**) Aggregation kinetics and the polymorphs outcomes like in **A** for the Era 1 (samples **45**-**4G**) and Era 2 (samples **53**-**62**) mixtures of full-length and NΔ4 HPLC-pure α-Syn. The time axes are all at the same scale for easier comparison and the darker shades of blue represent a higher percentage of NΔ4 as indicated in the figure.

Finally, across the Era 1 and Era 2 mixtures with NΔ4 it was clear that the pure NΔ4 α-Syn had a consistently longer lag-time (Figure 6B and 6C). One possible explanation is the stabilization of the intramolecular interactions of the N- and C- terminal regions of α-Syn that have been proposed to inhibit aggregation. The NΔ4 truncation has one less carboxylate (Asp2) and one additional amine relative to the full-length N-terminally acetylated α-Syn which we used in all other experiments. These changes significantly increase the positive charge of the N-terminal region and may therefore enhance long-range electrostatic interactions with the negatively charged C-terminal region. Such interactions are thought to stabilize autoinhibitory conformations of monomeric α- synuclein that exert a self-chaperoning effect and reduce aggregation propensity (Bertoncini et al. 2005). However, monomer–fibril interactions have also been proposed to involve electrostatic attraction between the positively charged N-terminus of monomeric α-synuclein and the negatively charged C-terminal tails exposed on the fibril surface (Kumari et al. 2021). Thus, the same charge redistribution that may enhance intramolecular self-chaperoning could also influence intermolecular monomer–fibril interactions during secondary nucleation and fibril growth. While the net effect of these competing processes is difficult to predict, it provides a possible explanation for how low levels of N-terminal truncation could nevertheless exert a disproportionate influence on polymorph selection, as observed in Figure 2C. Interestingly, a recent study reported that removal of the first four residues of α-Syn can accelerate amyloid formation, particularly under phase-separation conditions that promote condensate maturation and surface-dependent nucleation (Thrush et al. 2025). This contrasts with the longer lag time observed for pure NΔ4 α-Syn under our assay conditions, suggesting that the effect of this truncation is not intrinsically pro- or anti-aggregation, but depends on the aggregation pathway favored by the experimental environment. In our system, the increased positive charge at the N-terminus of NΔ4 may stabilize long-range interactions with the acidic C-terminal region and delay primary nucleation, whereas under condensate- or surface-dominated conditions the same truncation may instead enhance nucleation through altered interfacial behavior.

### Structural and symmetry variation with polymorph classes

The structural variation we observe falls into two conceptually distinct but related categories: variation in the core fold topology within a polymorph class, and variation in the protofilament interface between assembly variants of the same fold. The former is illustrated by the missing N-terminal strand in the type 2_-NS_ variants and the latter by the A, B, and D interface variants within type 1. Both levels of variation can occur simultaneously and independently within the same sample, and neither is fully predictable from the aggregation conditions.

Originally, the concept of numbered classes was developed in an attempt at bookkeeping for the quickly growing library of polymorphs. Along with the continuous accumulation of new classes, it has become apparent that the original classes are too broad to capture the variety of forms that α-Syn can achieve and would require sub-classes to better describe this variation. A prime example is the variety of type 1 folds which was already noted in our previous work (Figure 8 of (Frey et al. 2024)). Type 1 is possibly the most structurally variable of all the classified polymorphs, presenting the strongest case for a subclass nomenclature and until now the variety of type 1 folds only existed in comparisons between isolated samples. Here we have found that even within the same sample, multiple type 1 variants can coexist: the HPLC-pure full-length α-Syn sample **59** in Table 1 contains two nearly equally-represented variants of the type 1A fold (Figure 7A). The two other cases in which we have observed the coexistence of type 1 polymorphs (samples **45** and **60** Table1) represent interface or filament assembly variants that are derived from the same type 1 fold variant. The first of these, sample **45**, is an Era 1 HPLC- pure sample in which the type 1/7 fibril coexisted with the type 1A (Figure S4C and S4D) and the second case, sample **60** is the asymmetric 1D interface in the 5% NΔ4 coexisting with type 1m (Figure S4B). The type 1D fibril is also unusual in that it is both asymmetric and comprised of two filaments of opposite polarity.

**Figure 7.**
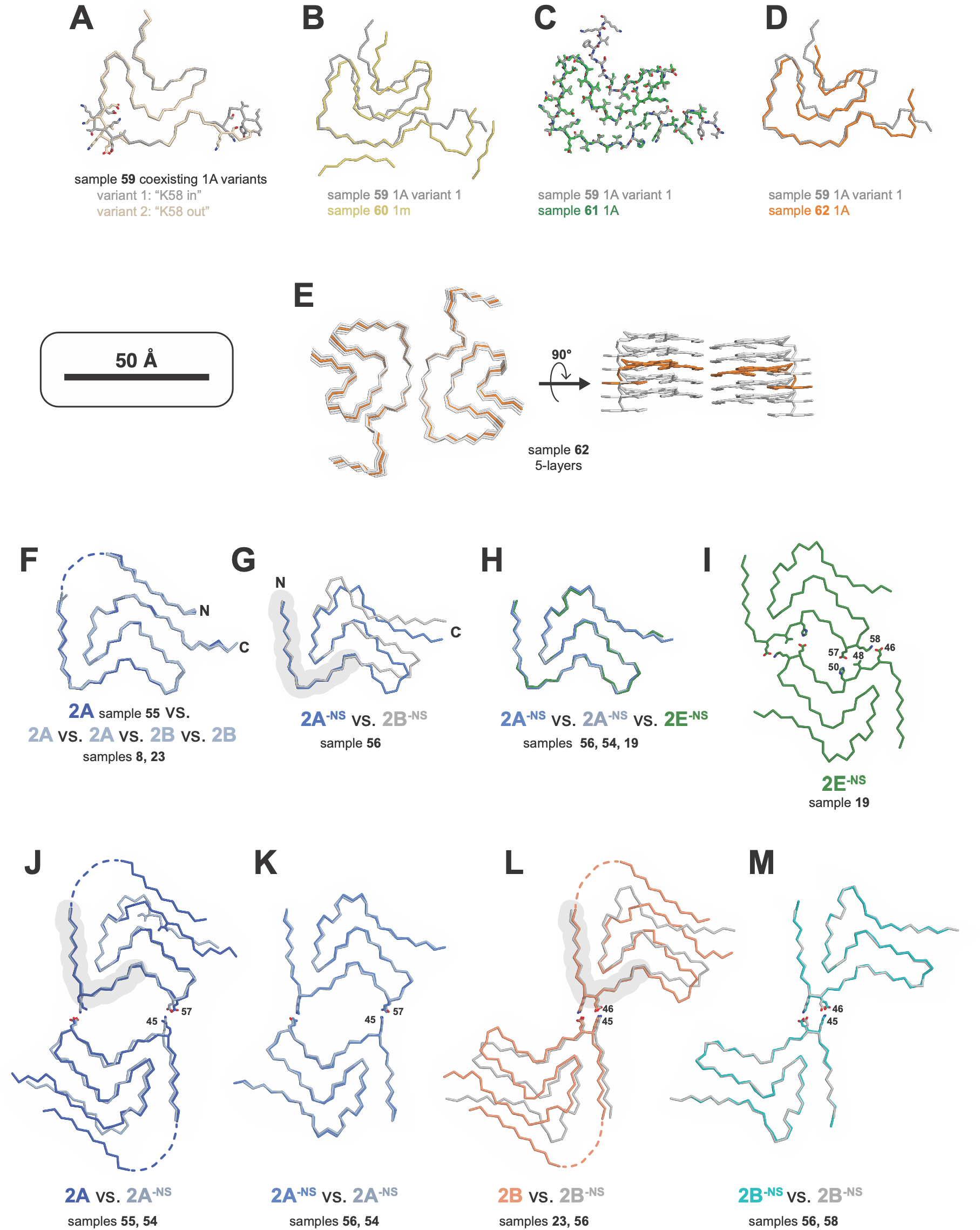
Structural variants within the types 1 and 2 polymorph classes. Overlay of type 1 structures from the same sample (**A**) and between samples (**B**)-(**D**) including the new “layer-traversing” fold (**E**) reveal significant variability in the type 1 fold within and between samples. The alignments of the type 1 structures was performed using all residues for **A** and **C** and for residues 50-63 for **B** and **D**. (**F**)-(**M**) Overlays of type 2^-NS^ and 2 structures (using alignments of residues 40-50) reveal fewer variations between three core folds. (**F**) Five independent type 2 structures from 2A and 2B fibrils are the same within and between samples. (**G**) The type 2^-NS^ structures are different between 2A^-NS^ and 2B^-NS^ fibrils derived from the same sample. (**H**) The type 2^-NS^ structures are the same between 2A^-NS^ and 2E^-NS^ fibrils. (**I**) The Cα trace of the novel 2E^-NS^ polymorph with the residues that make up its larger and more hydrophobic interface indicated. (**J**) Overlay of 2A and 2A^-NS^ structures shows their shared interface and the differences in the C-terminal portion of the fold with residue Ile88 highlighted to show its shifted but inward facing location in the last strand of the 2^-NS^ fold. The overlays were performed using only residues in the upper monomer for this and the remainder of the panels. (**K**) Overlay of 2A^-NS^ structures from different samples shows the lack of variability in this polymorph. (**L**) Overlay of 2B and 2B^-NS^ structures shows their shared interface and the differences angle between the 2^nd^ and 3^rd^ beta strands and in the C-terminal portion of the fold. (**M**) Overlay of 2B^-NS^ structures from different samples shows the lack of variability in this polymorph. The color of each Cα trace is paired with the sample or polymorph from which it is derived.

In another display of type 1 variety, there are five distinct type 1 polymorphs (one 1m, one 1D, and three 1A) identified among the four samples **59**-**62** which were all created from the same two stocks of α-Syn and NΔ4 (Figure 7 A-D). As mentioned above, the 1m and 1D have the same fold variant but both 1A folds from sample **59** and the single 1A fold in sample **62** are all different variants. Even the remaining 1A fold from sample **61** whose reconstruction reached 2.1 Å, while very similar to the “K58 in” variant from sample **59**, is still different in that its C-terminus is poorly defined in the density (Figure 7C, Figure S1). We attribute the unusually poor map quality at the high resolution of 2.1 Å to multiple distinct orientations and or twists that could not be resolved. Despite the large number of segments and high quality data of sample **61**, we were not able to separate the segments into different classes with distinct structures.

In contrast to the type 1 fold, the type 2 had been, in our hands and in the literature, relatively consistent. Now, and for no apparent reason, we have encountered a variety of type 2 structures. For example in the “era 2” HPLC-pure full-length and NΔ4 α-Syn mixed samples **54**-**58**, a type 2 that is missing the N-terminal beta strand (residues 14-25) was very well represented, dominating all except the 5% NΔ4 which had the typical types 2A and 2B with the N-terminal strand present. The fold of these type 2 variants that lack the N-terminal strand (-NS) 2A_-NS_ and 2B_-NS_ also diverge from the typical 2A/2B fold at other regions (Figure 7 G,J,L) but based on the inward facing orientation of the Ile88 sidechain, we still classify it as type 2. We are not the first to observe this type 2 variant as the 2B_-NS_ and a 2C_-NS_ have previously been reported as outcomes from a CSF-seeded aggregation (Fan et al. 2023). We encountered yet a new type 2 interface variant with the appearance of 2E_-NS_ in samples **19** and **20**, both of which were SP purity at pH 6.3 (Figure 7I). Still, the variation within the type 2 fold is tiny compared to type 1 with essentially only 3 main variants: the 2A and 2B folds are all very similar (Figure 7F), the 2A_-NS_ and 2E_-NS_ are nearly identical (Figure 7H), and lastly the 2B_-NS_ fold which is distinct from 2A/2B and 2A_-NS_/2E_-NS_ (Figure 7G).

In our large dataset of α-Syn structures a more subtle and perplexing variation in structure has also become apparent, namely a difference in the helical symmetry within the same interface variant. We have now observed this variation in two different polymorphs: the 3B and 5A can both exist in what appears to be C2 and pseudo-C2_1_ (2- start helical symmetry). We say “what appears to be” because in some cases, particularly with the 5A polymorph, segments can be aligned in relion_refine nearly as well in either symmetry, with both C2 and pseudo-C2_1_ giving contiguous density maps of superior quality to C1-based refinements (Figure S6). While in most cases the apparent symmetry can be visually determined in an “un-biased” C1-based refinement before any symmetry is applied or by visual inspection of high resolution 2D classes. In these straightforward cases, applying the correct symmetry during a 3D refinement will also yield a higher FSC resolution than with the wrong symmetry. However, in some cases the 2D classes display features that are consistent with both symmetries and with poor data, C1-based refinements do not always produce significant separation in the z-axis, precluding a proper helical symmetry assignment. In any case, it has been a mystery to us how maps of excellent quality can be produced in two different helical symmetries from the same data, a feat that is possible even in the less ambiguous cases in which the helical symmetry is visible in the rung patterns of the 2D classes and whose differences in FSC resolutions between the two symmetries is significant. Now we have encountered several high quality datasets with good z-separation whose symmetry remains ambiguous, often drifting between symmetries from one refinement to the next or one iteration to the next. Figure S6 highlights a case in which during C1-based refinement of 5A fibrils, the two half-maps appear to diverge into distinct symmetries, one more like C2 and the other more pseudo-C2_1_. We have tried to find any indications that the sample contains a mixture of fibrils with distinct helical symmetries, however we have had no success using 3D classification (with and without alignment) or 2D classification with filament subset selection to separate fibrils (or segments) of distinct helical symmetry. In the end, surprisingly subtle differences in map quality and perhaps a careful analysis of the contact distances at the dimer interface are the only clues that point to the correct symmetry. The warning here is that in some cases, the researcher should use extra caution before applying higher than C1 symmetry to avoid falling into a wrong refinement minimum.

### Structural Varia (Additional Structural Observations)

In the large amount of structural data presented in this manuscript there are many observations which are important but which do not add directly to the main findings. Here, for the record, we catalogue a few for the interested reader.

1. The large number of type 1m structures presented here reveal a high degree of similarity within this monofilament polymorph compared to the dimeric type 1A folds which has dozens of significant variants. Also, several of these type 1m reconstructions exceeded the resolution of our original deposition and the highest-resolution reconstruction presented (sample **15**) supports a revised sequence register for the residues in the N-terminal beta-strand whose EM density is detached from the rest of the fold. The original assignment of residues 13-20 to this density in our earlier model (PDB 8PK2) had relied on the positions of an analogous strand in a structurally similar monofilament fibril from juvenile-onset synucleinopathy (JOS) patient material (8BQV, (Yang et al. 2023). In our improved maps, an unambiguous assignment of the of residues 26-33 to a beta strand that packs against the strand formed by residues 36-41 was possible because the density for Gly31 clearly lacked a sidechain bump (Figure S7). Consequently, the original interpretation of the disconnected N-terminal strand was no longer viable due to the 27 Å distance that would need to be traversed by the newly placed residue Val26 and the previous end of the N-terminal strand Glu20. A shift of three residues in the N-terminal strand relative to our original interpretation (and to the JOS model) is the only possible interpretation of the density considering the sequence of the first 20 amino acids. Furthermore, in the revised model the electrostatic contacts are more sensible, with the sidechains of Lys10 and Lys12 instead of Glu13 being close to Glu57. Therefore, our current model PDB 32HW supersedes the coordinates in PDB 8PK2.
2. A striking feature that is readily apparent in some of the Cryo-EM reconstructions reported here (Figure S1) is the presence of density blobs of variable intensity that lie outside the protein fold at specific sites in several of the polymorphs. In certain datasets, these blobs are more intense (e.g. Type 2A sample **23**) while in others that are ostensibly from the same conditions they are less prominent (e.g. Type 2A sample **24**) and in others they are nearly absent (e.g. Type 2A sample **8**). Similar blobs are present to varying degrees in 1m and 5A and 5B reconstructions and always at the same location near surface lysine residues. Considering the aggregation conditions, the likely explanation is that phosphate from the buffer has become immobilized near these positively charged residues. While, it is not clear why there is such variability between datasets, it may be due to different degrees of order in the bound molecule or inherent differences in quality in the EM sample and/or data. These observations suggest that the charged surfaces of alpha-synuclein fibrils tend to attract whatever negatively charged molecules are present in the milieu and by extension, any observed density bound at the surface of fibrils might not have a physiological relevance.
3. Like the blobs in point 2 above, there are other features that are variable from one sample to the next within the same polymorph types. One example are the unidentified peptide chains that have only once been observed in a 5A structure (sample **52**) and are present in all 5m and 5B structures. A similar variable chain is present in the type 1/7 sample **45** but absent in the higher resolution map from sample **48** despite both samples coming from nearly identical buffer conditions. These minor differences in the amyloid fold could in principle have profound effects on the amyloid properties or biological activities due to the change in surface properties of the fibril.
4. The large variety of type 1 polymorphs in the literature makes it difficult to make sense of the conditions that lead to each unique fold. From an overview of deposited structures, it appears that certain labs can consistently produce the same Type 1A fold which suggests that there are still yet to be uncovered variables that can influence polymorph selection. With this in mind, the variety of type 1 structures that came from samples **59- 62** is particularly striking (Figure 7A-D) considering their common sample preparation and agitation protocol. This variety included a novel type 1 polymorph whose last two C-terminal strands traverse from one layer to the next within the amyloid fibril (Figure 7E). While layer-traversing amyloid folds are not new, to our knowledge this is the first case of an α-Syn amyloid making this type of intermolecular contact.

## CONCLUSION

One consistent finding from our analysis of nearly one hundred cryo-EM datasets of α- Syn fibrils is that no single variable is completely deterministic. Nevertheless, a coherent multivariable relationship between experimental conditions and polymorph outcome has emerged from this large collection of helical reconstructions. Variables that correlate with polymorph selection include pH, agitation conditions, sample purity, and the abundance of oligomers or seeds in the starting material. Together, these factors can explain much of the structural diversity reported in the literature and may account for some variation that has previously been attributed to post-translational modifications or disease-associated mutations.

Under gentle but continuous agitation and using HPLC-purified α-Syn preparations largely free of oligomeric species, pH becomes a major determinant of polymorph outcome, consistent with our previously described pH–polymorph correlation. In contrast, trace impurities such as the NΔ4 truncation can alter polymorph selection by suppressing formation of Type 5, a fold that incorporates a larger portion of the α-Syn sequence than any other polymorph reported to date and is uniquely characterized by involvement of the N-terminal methionine.

The influence of such trace impurities appears to be more pronounced in the Era 1 sample preparation protocol than in Era 2. We propose that this difference arises from a lower abundance of seeds or oligomers in Era 2 samples, thereby allowing pH to exert a stronger influence on polymorph selection. This interpretation is supported by the emergence of Type 5 as the dominant polymorph in Era 2 samples subjected only to 300 kDa filtration, whereas Type 5 likely represented only a minor species in the corresponding 100 kDa-filtered preparation.

The present study also expands the structural landscape accessible to wild-type full-length α-Syn fibrils through the identification of the new polymorphs 7, 8, and 9, found within the asymmetric heterodimeric fibrils 1/7 and 8/9, as well as a Type 1 interface variant in which two polar protofilaments adopt a head-to-tail arrangement within an asymmetric fibril. In addition, we observe previously undescribed structural variation within established polymorph classes, particularly Classes 1 and 2.

Collectively, these observations suggest that many α-Syn polymorphs occupy a relatively narrow thermodynamic landscape and that kinetic effects play a major role in determining which structure ultimately emerges. Small differences in solution conditions, impurity profiles, or the presence of oligomeric species can therefore influence fibril formation toward distinct structural outcomes.

The present results also have practical implications for reproducibility across laboratories. Protein purity, the method used to prepare the monomeric starting material, and the agitation hardware and protocol are rarely standardized or fully reported in published aggregation studies, yet each of these variables demonstrably alters polymorph outcomes. Additionally, the unexpected N-terminal methylation of the NΔ4 construct, observed in two independent expressions and undetectable by SDS-PAGE, serves as a reminder that chemical heterogeneity in recombinant α-Syn preparations is not limited to truncation products. Laboratories that do not co-express NatB and therefore produce α-Syn with a free N-terminal amine may also carry uncharacterized N-terminal methylation, which could influence aggregation behavior in ways that have not been appreciated

Although the original goal of reproducing the disease-relevant Parkinson’s disease polymorph in vitro remains unmet, we believe that the present work provides important insight into the origins of α-Syn polymorph diversity and the experimental variables that govern its formation. A better understanding of these relationships should improve the reproducibility of *in vitro* fibril preparations and facilitate the rational design of polymorph-specific probes, inhibitors, and therapeutic strategies for the study and treatment of synucleinopathies.

## Materials and Methods

### Recombinant protein expression and purification of α-Syn

Recombinant N-terminally acetylated, human WT α-Syn was produced by co-expression with the yeast N-acetyltransferase complex B (NatB) in the (Johnson et al. 2010) plasmid in Escherichia coli BL21 Star™ (DE3) (Thermo). α-Syn was expressed and purified according to the previously published protocol (Campioni et al. 2014) with slight modification. In brief, a single colony from a double transformation with the NatB and α-Syn plasmids was grown at 37°C in 100 ml LB media containing 100 µg/ml ampicillin and 50 µg/ml chloramphenicol to an OD of 2 and stored at -80° C after the addition of 30% glycerol. A scrape from the glycerol stock was used to inoculate an overnight primary growth culture with the same antibiotic selection which on the next day was diluted 40-fold into freshly prepared LB media again with the same antibiotics. Protein over-expression was initiated by the addition of 1 mM IPTG when the OD reached 0.8 and after 4 hours of induction at 37°C the cells were harvested by centrifugation at 5000 g and stored frozen at -20 °C. The purification of α-Syn from the periplasmic fraction closely followed a previously published protocol (Huang et al. 2005). The frozen pellet was thawed and resuspended in 65 ml of osmotic shock buffer (30 mM Tris pH 7.2, 40% sucrose, 2 mM EDTA) per liter of original cell culture, incubated at room temperature for 10 min and then pelleted at 8000 g for 15 min at 25° C. The cell pellet was resuspended in 65 ml cold H2O followed immediately by the addition of 37.5 µl saturated MgCl2 and incubation on ice for 3 minutes. Cells were again pelleted at 8000 g for 20 min at 10° C, 8000 g and the supernatant stored at -25 ◦C. The periplasmic fraction was loaded onto a HiTrap Q FF column (Cytiva), equilibrated with 20 mM Tris, pH 7.0. The bound protein was eluted using a gradient to 20 mM Tris pH 8.0, 750 mM NaCl. Solid [NH_4_]_2_SO_4_ was then added to the α-Syn containing fractions to a final salt concentration of 1 M. The sample was loaded onto a HiPrep Phenyl FF 16/10 column (Cytiva), equilibrated with 50 mM NaPO_4_ pH 7.0, containing 1 M [NH_4_]_2_SO_4_. After washing with the equilibration buffer, α-Syn was eluted in a gradient against 50 mM sodium phosphate without [NH_4_]_2_SO_4_. The eluted protein was dialyzed extensively against water with a 6-8 kD MWCO membrane (SpectraPor®, Repligen), frozen in aliquots in liquid nitrogen, lyophilized and stored at –20° C as standard purity (SP) protein. The SP protein was further purified by resuspending the lyophilized samples at 10-15 mg/ml in 10% CH_3_CN, 0.1% TFA, loading about 2-4 mg onto a semi-prep (10 x 250mm) Jupiter 5 µm C4 300A reverse phase column (Phenomenex) pre-equilibrated in 10% CH_3_CN, 0.1% TFA and eluting the protein in a gradient of CH_3_CN, 0.1% TFA. The HPLC purified fractions were pooled, frozen in aliquots in liquid nitrogen, lyophilized and stored at –20° C.

### Monomer sample preparation for aggregation experiments

Two different protocols were used to prepare soluble α-Syn. The first, referred to as “Era 1” started with dissolving the lyophilized α-Syn in PBS (P4417, Sigma-Aldrich) at around 500 µM. The pH of the dissolved α-Syn was adjusted to pH 7.5 and then passed over a PD-10 column pre-equilibrated in PBS at the pH of interest. The buffer-exchanged protein was concentrated back to about 500-600 µM using an Amicon Ultra-4 10 kD MWCO device (Millipore) and then passed over a VivaSpin 500 filter with a 100 kD MWCO (Sartorius) to remove aggregates. The final filtering caused a loss of 30-50% of the sample. The final concentration was measured and the sample diluted to the desired concentration (typically 300 µM) with 0.2 µm filtered buffer. The second protocol, referred to as “Era 2” started with dissolving the lyophilized α-Syn in pure H2O. This procedure was designed around the HPLC-purified α-Syn that when dissolved at high concentration in water gave slightly acidic solutions due to the trace TFA from the HPLC solvents. The non-buffered protein could be solubilized at 700-800 µM and then then dialyzed in less than 500 µl volumes using a homemade device comprising the cap of a 5 ml Eppendorf tube with a piece of a 6-8 kD MWCO membrane held in place by a ring obtained from the top 2 mm of the body of the 5 ml tube. After 48 h of dialysis against a 1000 fold volume of the sample with one buffer change after 24 h, the sample was recovered by piercing the membrane with a needle and then filtered over a series of VivaSpin 500 devices: first with a 300 kD MWCO and then with a 100 kD. The filtering in series prevented fouling of the 100 kD membrane thereby speeding up the process and also bringing down the total loss of sample from the initial solubilization to less than 10%. The protein was then diluted with 0.2 μm-filtered buffer to the desired concentration.

### α-Syn aggregation

Two main protocols were used in this study. The first “rotation” method was to put 500 µl of the “Era 1” protein solution in a 1.5 ml Eppendorf Protein LoBind tube and then incubate it with gentle agitation via tumbling (∼20 rpm) at 37°C for 5 days. The second “plate reader” method was carried out in two different types of 96-well plates: µCLEAR® non-binding (Greiner P/N 655906) and UV-STAR® (Grein P/N 655809) in a Pherstar (BMG) platereader. The µCLEAR plates were adopted so that we could also monitor the tyrosine fluorescence polarization (FP) during the aggregation, however they lack the hydrophilic coating that should reduce non-specific binding. The aggregation was carried out with 200 µl of sample per well with intermittent 250 rpm orbital shaking as part of the aggregation kinetics measurement protocol. The measurement cycle involved 30 s shaking before the measurement of absorbance at 340 nm (ca. 1 min) followed by 30 s shaking before the FP measurement (ca. 1 min) followed by 15 min of gentle movement of the plate inside the plate reader to minimize thermal gradients.

### Electron microscopy grid preparation and data collection

Cu R2/2 300 mesh grids (Quantifoil) were glow-discharged at 25 mA for 30 s and 120 s, respectively. Freshly glow-discharged grids were used in a Vitrobot Mark IV (Thermo Fisher Scientific) with its chamber set at 100% humidity and at a temperature of 22°C. Fibrils (4 µl aliquots) were applied to the grid and blotted for 8–15 s after a 30–60 s wait time, and subsequently plunge-frozen into a liquid ethane/propane mix. The grids were clipped and immediately used or stored in liquid nitrogen.

Data acquisition was performed on a Titan Krios (Thermo Fisher Scientific) operating at 300 kV equipped with a Gatan Imaging Filter (GIF) with a 20 eV energy slit using Gatan’s K3 direct electron detector in counting mode. Movies were collected using EPU software (Thermo Fisher Scientific) at a magnification of ×130k and a dose rate of approximately 8 e/pixel/s and total dose ca. 62–74 e/Å_2_.

### Image processing

Image processing and helical reconstruction were carried out with RELION 5.0, following the procedure for amyloid structures as described in (Scheres 2020). Movies were gain-corrected and RELION’s own motion correction was used to correct for drift and dose-weighting and CTF estimation was done using Ctffind4.1 (Rohou and Grigorieff 2015). The exact details of processing and refining the structures from each dataset varied but followed a similar pipeline as outlined here. Individual fibrils were manually selected from a subset of the ca. 100 micrographs and the extracted segments (333 Å box, 2.6 Å/px) were used to get initial 2D classes. The segments from better classes were used to train autopicking with the Topaz implementation in RELION. The segments from autopicking were extracted as before and used for 2D classification with 300-1000 classes depending on the number of segments, aiming to keep the ratio of segments to classes below 1000. The segments from different polymorph classes were separated using the either the standalone FilamentTools (David Li), a collection of scripts for processing cryo-EM filament data available through GitHub or its implementation in RELION 5.0 (Lovestam et al. 2024). An initial model was generated from the separated 2D classes with relion_helix_inimodel2d. After an initial alignment in Refine3D which typically reached the Nyquist limit of 5.2 Å, the refined particles were re-extracted at either 1.3 Å/px in a smaller 260 Å box (with the exception of wider 5B polymorphs and sample **6** and **7** from Table 1 which were kept at the original 330 Å box size). Bayesian polishing (Zivanov et al. 2019) and CTF refinement (Zivanov et al. 2020) were performed with intermittent 3D refinements to improve the resolution of the reconstructions. The maps of the final reconstructions were sharpened using standard postprocessing procedures in RELION.

### Model building and refinement

Initial protein coordinates were built into the postprocessed maps with ModelAngelo (Jamali et al. 2024), the output of which was manually adjusted in COOT (Emsley and Cowtan 2004) and further improved by real-space refinement as a nine-layer fibril in ISOLDE (Croll 2018) with symmetry restraints. The coordinates of the 9-layer fibril were further refined in PHENIX (Liebschner et al. 2019) to obtain reasonable b-factors after which the outer two layers at each end of the fibril, which often diverge slightly in structure due to their placement at the edges of the model, were removed for deposition of the central five layers in the PDB. Figures were prepared with CCP4MG (McNicholas et al. 2011).

### Proteomics sample preparation, data acquisition and analysis

Enzymatic digestion with endopeptidase GluC was performed on 10 μg of protein in 10 mM Tris pH 8 with 2 mM CaCl_2_. After digestion the samples were dried and then dissolved in aqueous 3% Acetonitrile with 0.1% formic acid. The peptide concentration was estimated with the Lunatic UV/Vis absorbance spectrometer (Unchained Labs) before separation on an M-class UPLC and analyzed on a Orbitrap mass spectrometer (Thermo). The MS data were processed for identification using Byonic 5.10 (Protein Metrics, USA). The spectra were searched against the *E. Coli* database plus the NΔ4 α-Syn sequence including the following modifications: Oxidation (Met), Methylation (Protein N-terminus), Methylation (at any amino acid)

## Acknowledgements

The authors gratefully acknowledge the Functional Genomics Center Zurich (FGCZ) of University of Zurich and ETH Zurich, and in particular Dr. Chia-Wei Tan-Lin, for the support on the proteomics analyses. Financial support was obtained from the Synapsis foundation, grant number 2023-PI04.

---

**Figure S1.**
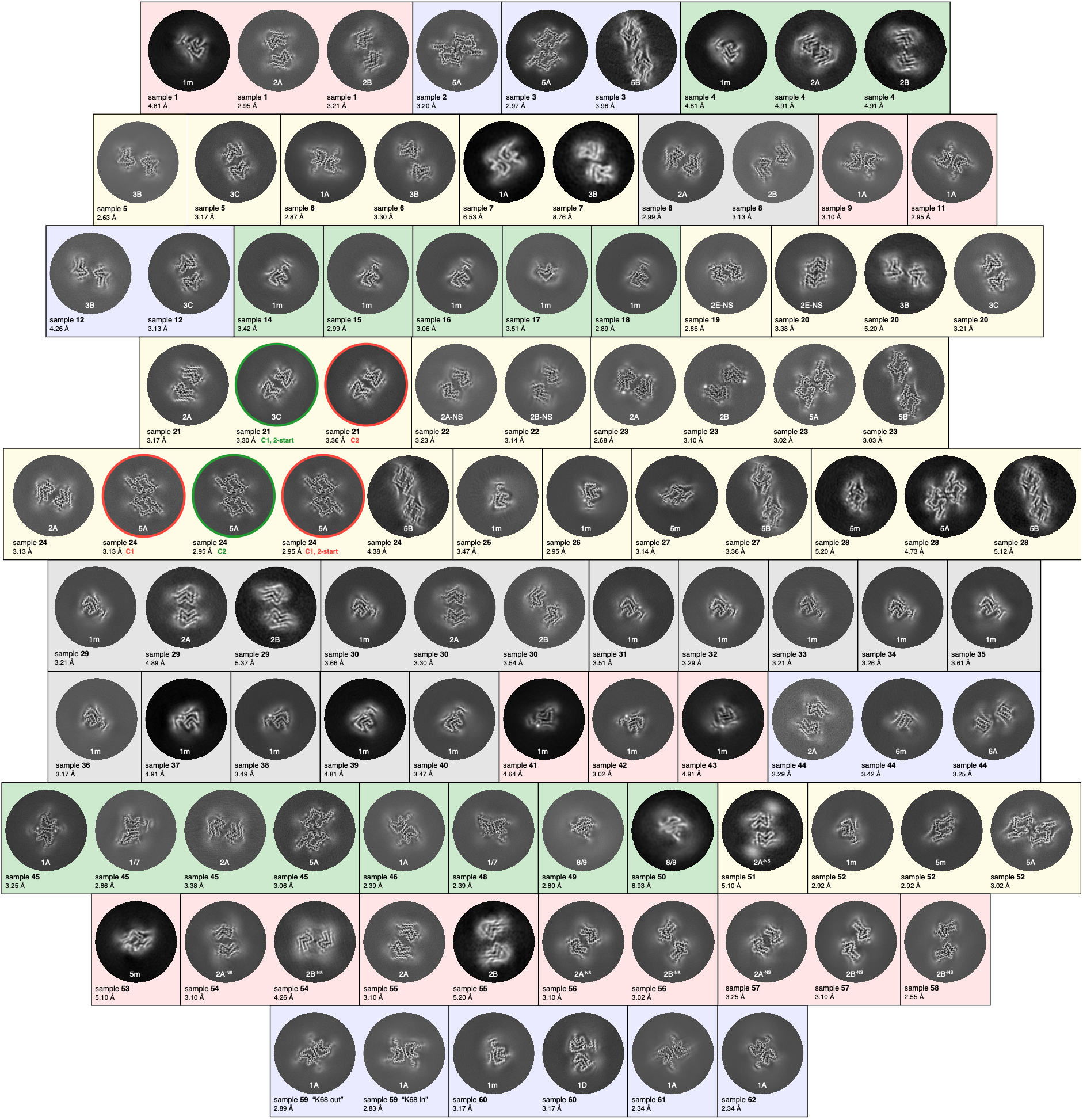
Overview of all helical reconstructions reported in this work. The sample numbers refer to the listing in Table 1 and the images show an approximately a 5 Å thick slice of the cryo-EM map in a 200 Å diameter region that is centered on the helical axis. The contiguous background colors connect samples that were prepared and aggregated at the same time while the black outlines indicate individual samples, some of which contain multiple helical reconstructions. The two samples with red and green circles around the z-slices are to highlight the two examples discussed in the manuscript for which the helical symmetry was somewhat ambiguous. The green circles are around the reconstruction that gave the best resolution/or map quality. The reported FSC resolutions are those before postprocessing in RELION.

**Figure S2.**
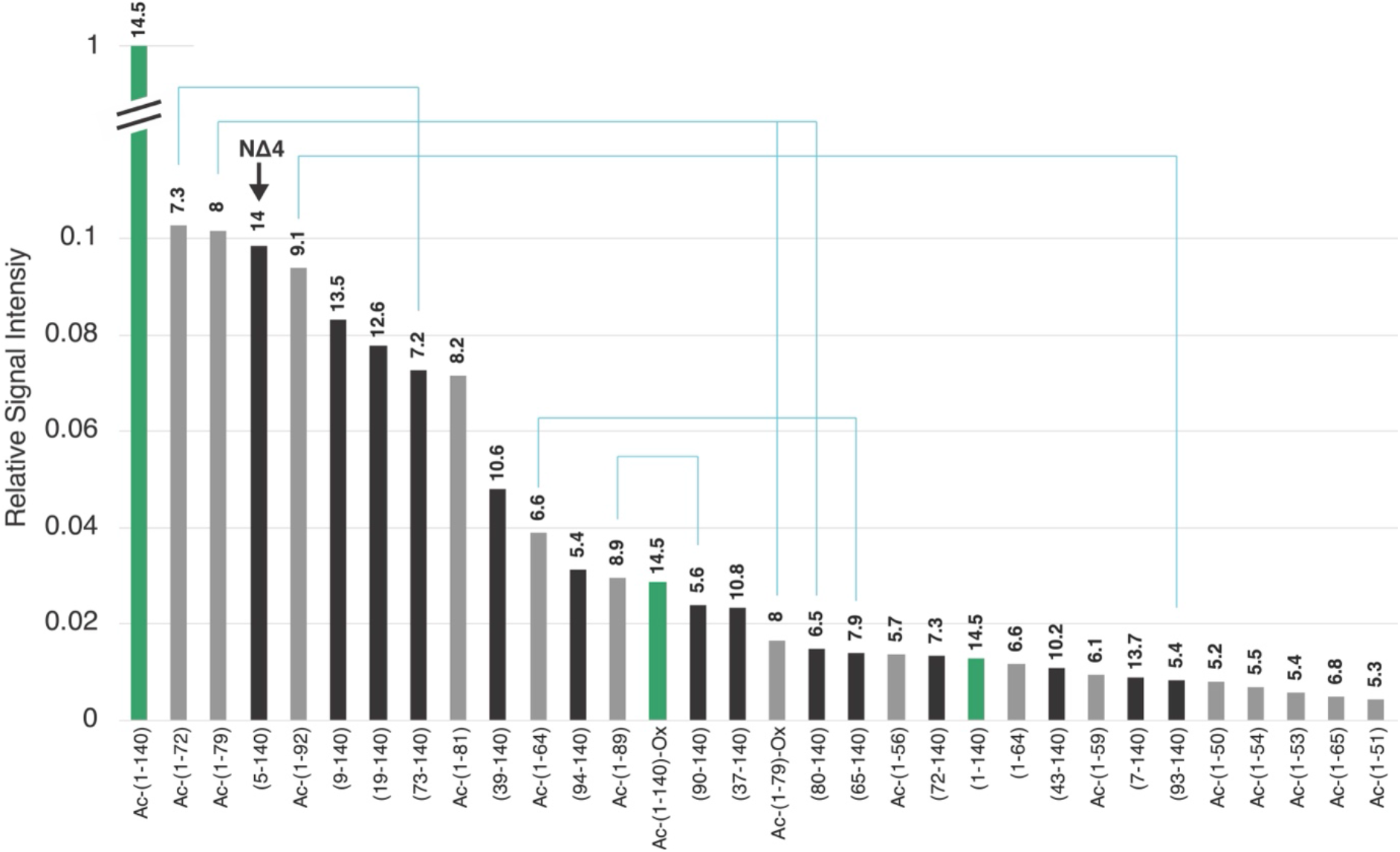
Peak intensities of aSyn and its truncation products identified by LC-MS qTOF analysis. The 25k g pellet fraction of a SP-purity aSyn aggregation that produced type 1m fibrils was solubilized in 6M guanidine and then injected on an Agilent Eclipse Plus C18 column and eluted with a gradient of CH3CN with formic acid as an ion pairing agent for direct analysis on a Bruker Daltonics maXis ESI-QTOF instrument. The peaks whose mass fell within 5 ppm of the calculated mass of an aSyn fragment (the largest or second largest peak of a calculated isotope envelope) were tabulated according to their relative intensity. All combinations of acetylated, oxidized and unmodified masses were searched. The green bars represent the full-length aSyn and the black and grey are the N- and C-terminal truncations. The mass of each peak in kD are listed above each bar. The light blue lines connect fragments that are formed from a single cleavage of the full-length protein. Peaks that likely represent protein with an oxidized Methionine are noted with “-Ox”

**Figure S3.**
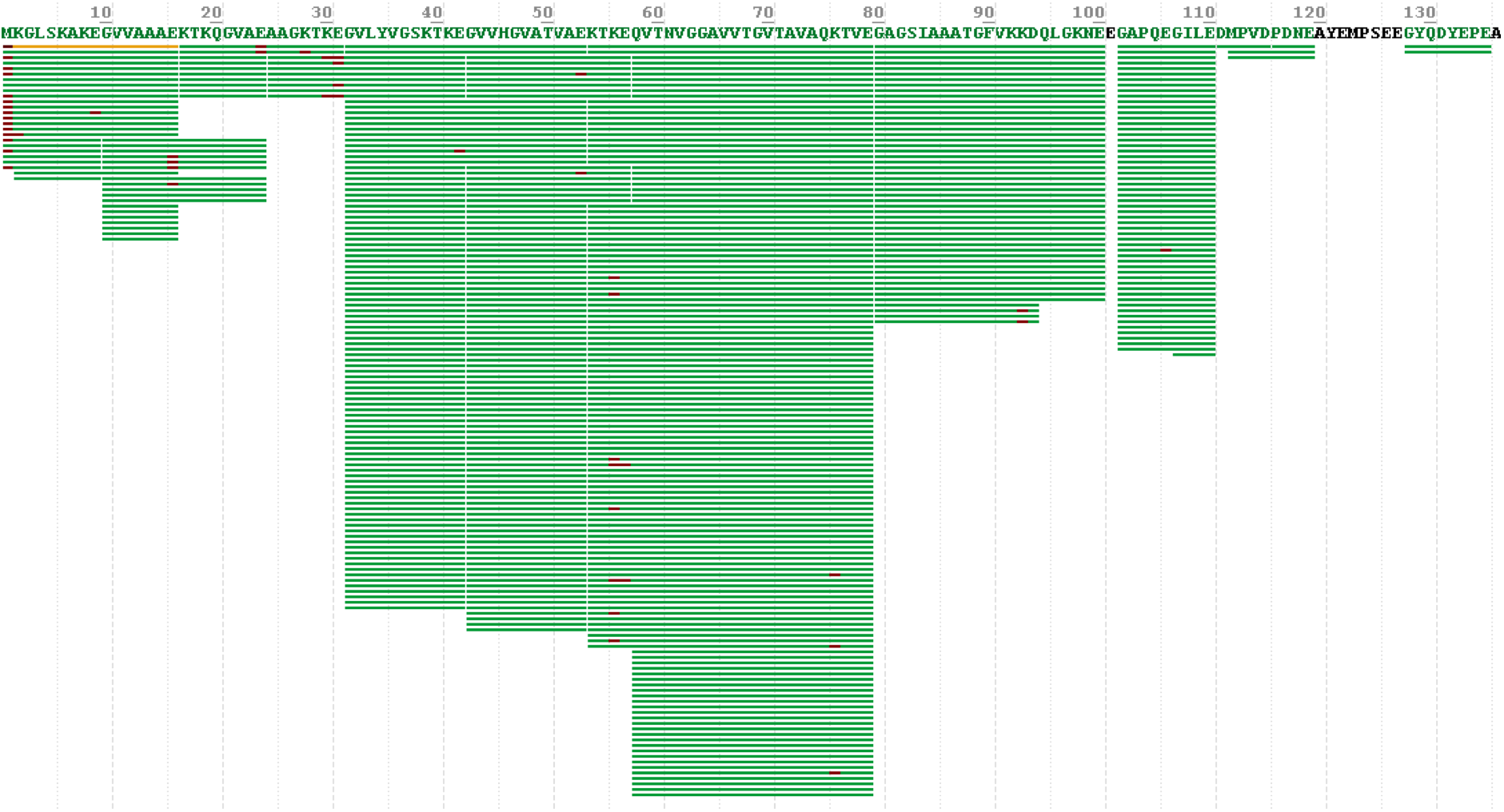
Sequence coverage of α-Syn endopeptidase GluC peptides in NΔ4 sample. The green bars represent peptides identified by LC–MS/MS analysis as having an extra CH2 mass, while the amino acid highlighted in red indicates the modified residue.

**Figure S4.**
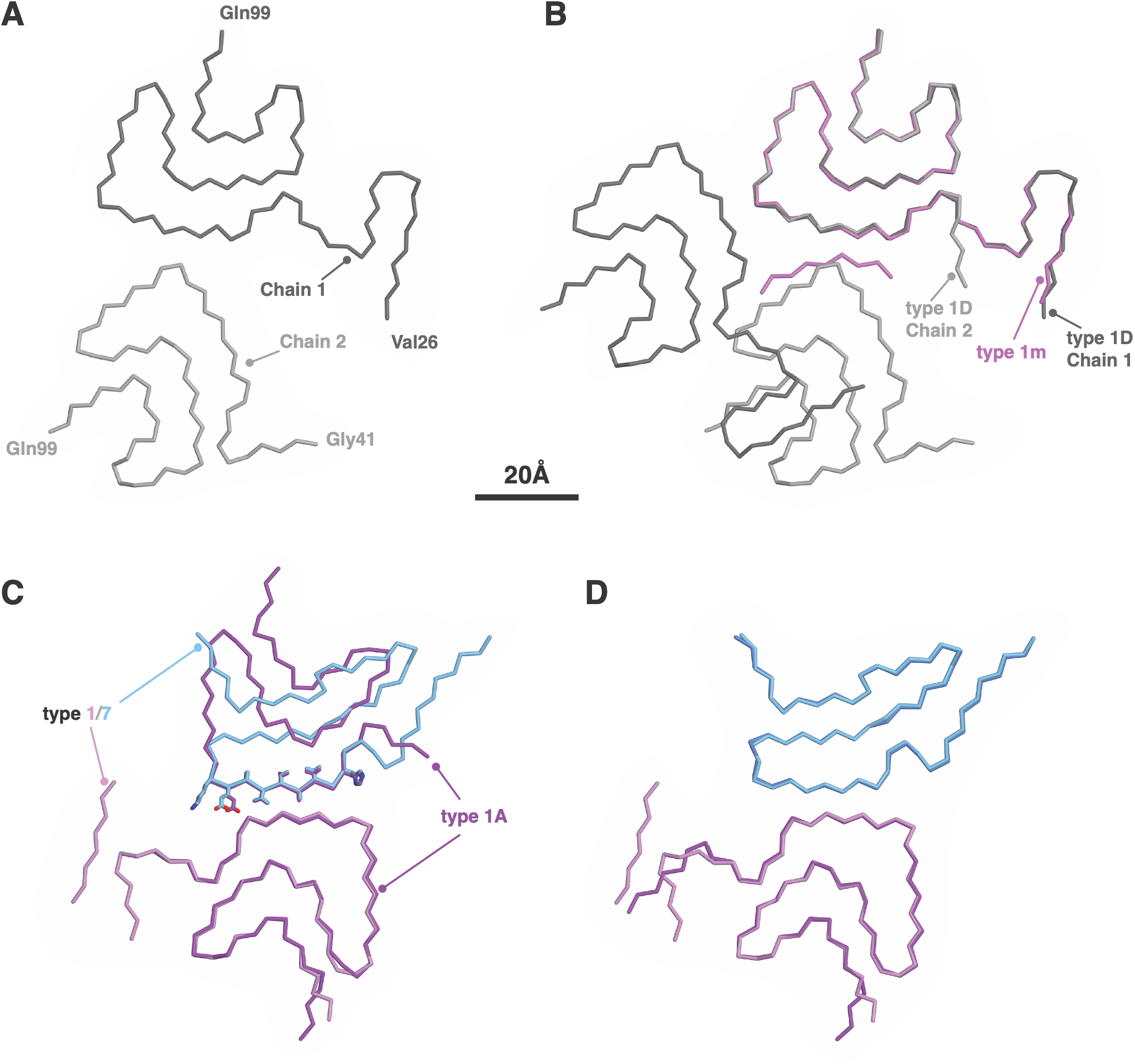
Structural conservation of type 1 fold within coexisting interface variants. (**A**) The type 1D fold from sample **60** is shown as a CA trace with its two chains lighter and darker shades of grey. (**B**) The structural similarity of the type 1m and 1D coexisting polymorphs is shown in an overlay of the three chains (chains 1 and 2 from type 1D with 1m). The 1D chains are colored as in **A** and the 1m chain is in light purple. (**C**) The structural similarity of the type 1A and 1/7 coexisting polymorphs in sample **45** is shown in an overlay with the 1A chains in darker purple and the two chains from 1/7 colored light purple (type 1) and light blue (type 7). The alignment of the structure was performed using only residues 40-90 of the type 1 chain from the 1/7 polymorph. The sidechains for residues 50-58 are shown in the type 7 chain and the type 1 chain with which it coincides in order to show the similarity between the two interfaces. (**D**) The type 1/7 polymorph from sample 45 (colored as in **C**) is overlayed with that of sample **48** (in darker shades of purple and blue) to show the conservation of the core fold with the variability in the N and C termini of the type 1 chain.

**Figure S5.**
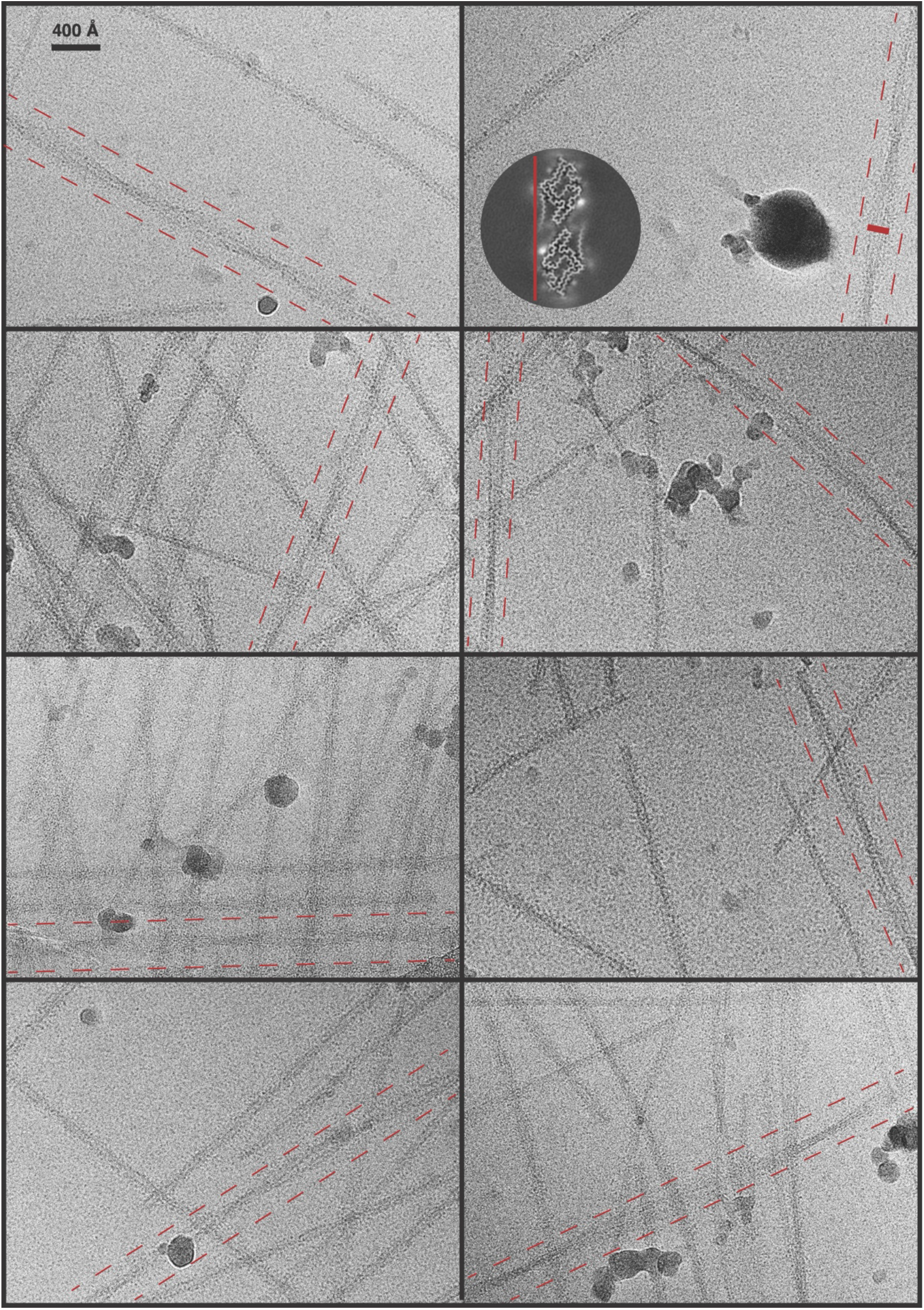
Fibrils with 5B-like morphology. Eight selected micrographs with fibrils that display a 5B-like morphology. The unique cross-section of the 5B polymorph makes its identification in micrographs relatively easy and the suspected 5B fibrils are highlighted by maroon dotted lines on each side. The widest section of a 5B-like fibril in the upper right micrograph is overlayed with a maroon scale bar representing 180 Å, matching the end-end longest dimension of the 5B dimer. The z-slice of a 5B map from another dataset is overlaid as a reference with the vertical maroon line also scaled to 180 Å.

**Figure S6.**
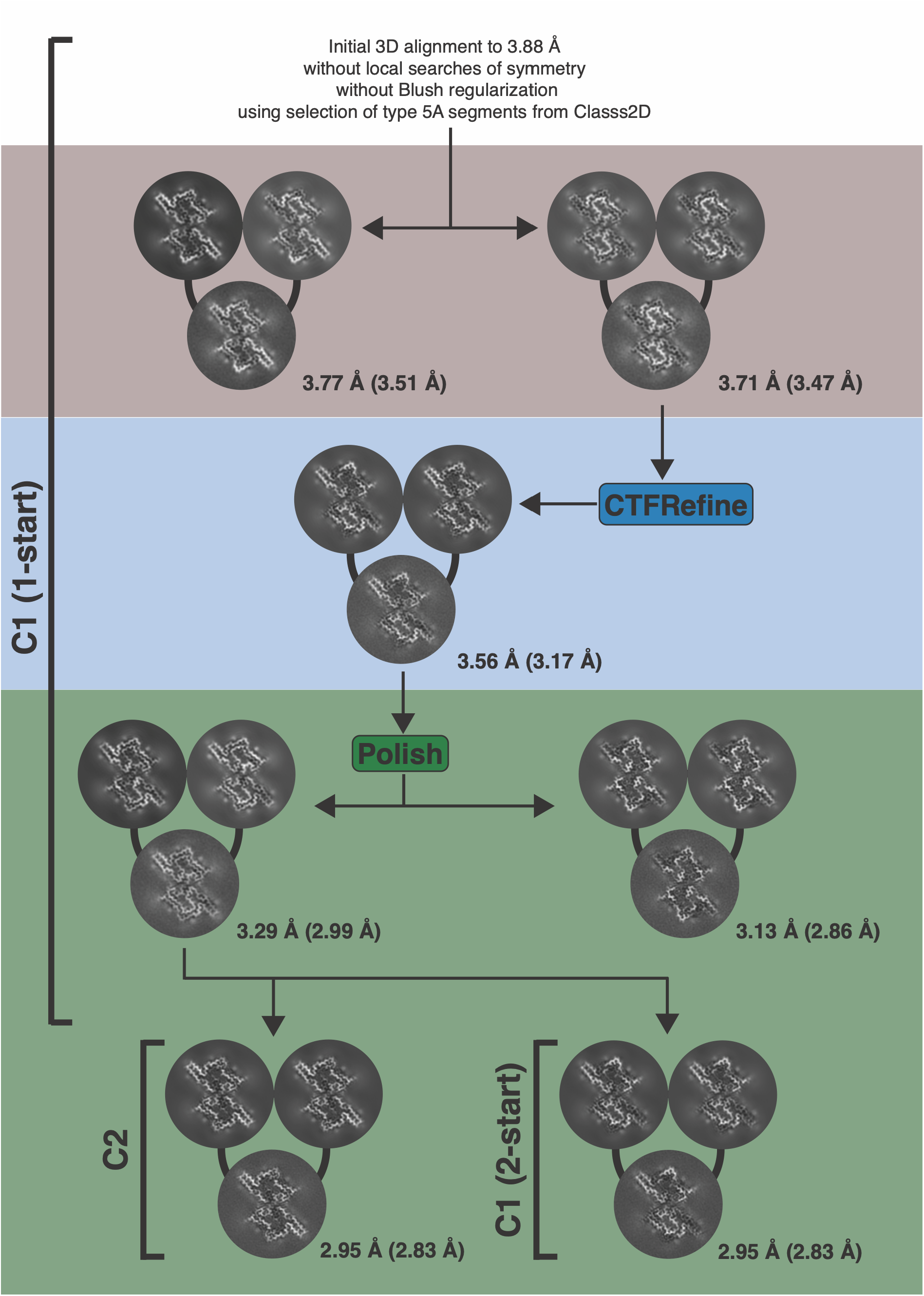
Refinement trajectory for a single set of type 5A helical segments. The ambiguous helical symmetry of sample **24** is highlighted with each set of three images showing the central layer of the final half-maps and the combined final map from a Refind3D job in RELION. The arrows indicate the jobs from which input data where derived. The helical symmetry that was applied during each reconstruction is indicated by the brackets to the left of the images with C1 being used exclusively until after the Bayesian polishing step. The FSC resolutions before and, in parentheses, after postprocessing are indicated. While the best map appeared for the C2 symmetry, its improvement over that of the 2-start symmetry was not large, and the half-maps during the C1 refinements sometimes displayed more of a 2-start symmetry than a C2 symmetry. Even the half-maps with a single iteration did not have the same apparent symmetry (top left job) and these disagreements between half-maps was variable from iteration to iteration and job to job. For example, after the Polish job, the two identical refinements run in C1 led first to a 2-start-like output map (left) and then later to a clear C2 symmetry (right), the latter having a higher FSC resolution. The output of the former of these two jobs was used in both C2 and 2-start helical refinements that each ended with the same final FSC resolution. The refinement with the applied C2 symmetry gave a slightly improved map quality and so this was chosen as the “correct” symmetry for this dataset.

**Figure S7.**
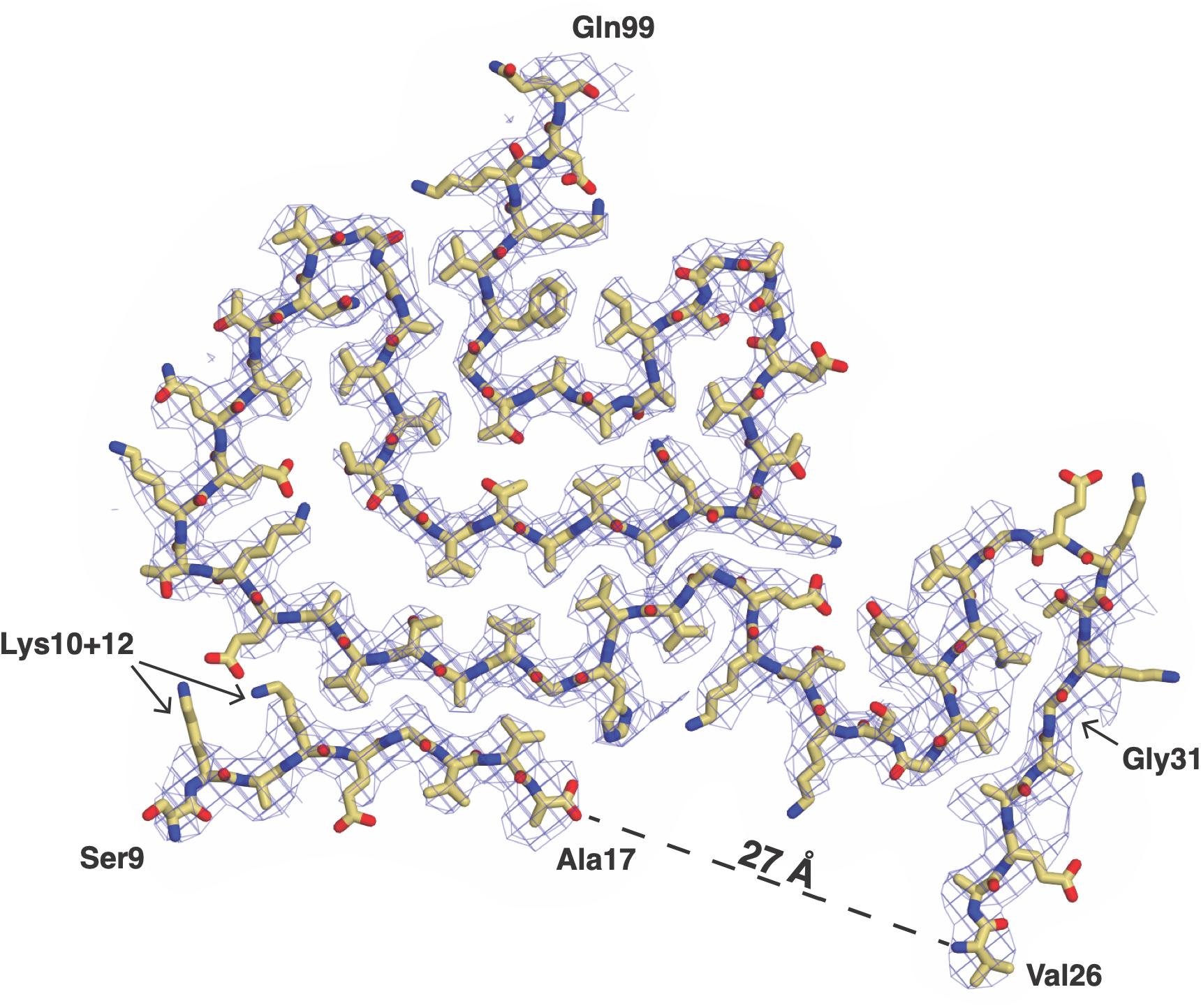
The revised model of type 1m from sample **15** overlayed with its cryo-EM density contoured at 4σ. The termini of the modeled chains are labelled as well as the positions of Lys10, Lys12 and Gly31 as discussed in the text.

**Table S1:** PDB and EMDB entry codes. The dataset which have been deposited in the PBD and EMDB databases are listed. _1_The data from the 5A polymorphs was reconstructed under three different helical symmetries as illustrated in Figure S6. All three maps have been deposited as an example of ambiguous symmetry. _2_The information here is redundant with the main text and Table 1 but is included to help orient the reader to the unique features of certain structures.

| Sample | Polymorph | PDB entry | EMDB entry | Notes<sup>2</sup> |
| --- | --- | --- | --- | --- |
| 12 | 3C | 32HY | 58905 | C1, 2-start |
| 15 | 1m | 32HW | 58903 |  |
| 19 | 2E<sup>-NS</sup> | 32IG | 58913 |  |
| 23 | 2A | 32DS | 58821 |  |
| 23 | 2B | 32ED | 58830 |  |
| 23 | 5A | 32EP | 58842 | C2 |
| 23 | 5B | 32ER | 58845 |  |
| 24 | 5A<sup>1</sup> | - | 58899 | C2 |
| 24 | 5A<sup>1</sup> | - | 58900 | C1, 2-start |
| 24 | 5A<sup>1</sup> | - | 58901 | C1 |
| 44 | 6m | 32IR | 58924 |  |
| 44 | 6B | 32IS | 58925 |  |
| 45 | 1A | 32IH | 58914 |  |
| 45 | 1/7 | 32IK | 58917 |  |
| 45 | 2A | 32IL | 58918 |  |
| 45 | 5A | 32IM | 58919 | C1, 2-start |
| 46 | 1A | 32IO | 58921 |  |
| 48 | 1/7 | 32IP | 58922 |  |
| 49 | 8/9 | 32IQ | 58923 |  |
| 56 | 2A<sup>-NS</sup> | 32IE | 58911 |  |
| 58 | 2B<sup>-NS</sup> | 32IF | 58912 |  |
| 59 | 1A | 32HZ | 58906 | 1 of 2 co-existing variants |
| 59 | 1A | 32IA | 58907 | 2 of 2 co-existing variants |
| 60 | 1D | 32IB | 58908 |  |
| 60 | 1m | 32IS | 58926 |  |
| 61 | 1A | - | 58927 | choppy density at 2.1 Å |
| 62 | 1A | 32IC | 58909 | "layer traversing" variant |

